# Bacterial type 1A topoisomerases maintain the stability of the genome by preventing and dealing with R-loop-and nucleotide excision repair-dependent topological stress

**DOI:** 10.1101/2021.07.10.451908

**Authors:** Julien Brochu, Emilie Vlachos-Breton, Marc Drolet

## Abstract

*E. coli* type 1A topoisomerases (topos), topo I (*topA*) and topo III (*topB*) have both relaxation and decatenation activities. *B. subtilis* and *E. coli topA topB* null cells can survive owing to DNA amplifications allowing overproduction of topo IV, the main cellular decatenase that can also relax supercoiling. We show that overproducing human topo IB, a relaxase but not a decatenase, can substitute for topo IV in allowing *E. coli topA* null but not *topA topB* null cells to survive. Deleting *topB* exacerbates phenotypes of *topA* null mutants including hypernegative supercoiling, R-loop formation, and RNase HI-sensitive replication, phenotypes that are not corrected by topo IV overproduction. These phenotypes lead to Ter DNA amplification causing a chromosome segregation defect that is corrected by topo IV overproduction. Furthermore, *topA topB* null mutants not overproducing topo IV acquire *uvrB* or *uvrC* mutations, revealing a nucleotide excision repair (NER)-dependent problem with replication fork progression. Thus, type IA topos maintain the stability of the genome by providing essential relaxase and decatenase activities to prevent and solve topological stress related to R-loops and NER. Moreover, excess R-loop formation is well tolerated in cells that have enough topoisomerase activity to support the subsequent replication-related topological stress.

## INTRODUCTION

Because of the double helical structure of DNA, each time the two strands are separated during replication, transcription, or repair, underwinding and overwinding (supercoiling) occur. In turn, such supercoiling interferes with normal gene expression and replication. Furthermore, tangling of the DNA occurs during replication and repair. If not properly resolved, such entanglements inhibit chromosome segregation and may lead to DNA breaks and genomic instability. To solve these topological problems, cells possess DNA topoisomerases (topos), which are nicking-closing enzymes (1,2). These enzymes cut one (type I) or two (type II) DNA strands. Type I and II are further divided respectively in class IA and IB and IIA and IIB. To solve topological problems, type IA and type II enzymes use a strand-passage mechanism whereas enzymes of the class IB family, also named swivelases, use a rotation mechanism.

Class A enzymes are the only ubiquitous topos (3,4). Furthermore, unlike other topos, they use ssDNA as substrates and many of them possess RNA topo activity (5,6). Type IA topos are classified into three subfamilies, the two main ones being topo I and topo III with *E. coli topA* and *topB* respectively encoding the prototype enzyme of these two subfamilies. The third one, reverse gyrase, is only found in hyperthermophilic and some thermophilic organisms. Topo I is present in all bacteria but not in archaea and eukaryotes, whereas topo III is found in most bacteria and in all archaea and eukaryotes. Topo III has a higher requirement for ssDNA than topo I and mostly acts as a decatenase (7), whereas topo I mostly acts to relax transcription-induced negative supercoiling (8). Notwithstanding these preferences, it is quite clear from *in vitro* experiments that topo I and III can perform both relaxation and decatenation reactions (9–11). The results of recent single-molecule experiments suggest that a dynamic fast gate for topo I may promote efficient relaxation of negatively supercoiled DNA, whereas a slower gate-closing for topo III may facilitate capture of dsDNA and, as a result, efficient decatenation (12). Additional factors that must be considered for their *in vivo* activity are, among others, their relative abundance which is reported to be about 200 and 20 ppm respectively for topo I and topo III (https://pax-db.org/species/511145), and their interacting partner, e.g. RNAP for topo I (13,14) and the replication fork for topo III (likely via SSB and DnaX; ref. (15,16)).

Null mutations in the *topA* gene of *E. coli* inhibit cell growth. Few *topA* null cells manage to generate visible colonies owing to the acquisition of compensatory mutations (reviewed in (4)). Such mutations are either substitutions in *gyrA* or *gyrB* genes that reduce the negative supercoiling activity of gyrase (17,18), or amplification of a chromosomal region allowing topo IV overproduction (4,19–22). The occurrence of gyrase mutations is easy to interpret since the *in vivo* function of gyrase is the introduction of negative supercoiling (via the relaxation of positive supercoiling generated during replication and transcription). However, the main function of topo IV is to act as a decatenase behind the replication fork to remove pre-catenanes and, once chromosomal replication is completed, to remove catenanes to allow chromosome segregation (23,24). Topo IV can also relax negative supercoiling, albeit much less efficiently than positive supercoiling relaxation and decatenation (25–27). Because of the established main function of topo I, it is believed that topo IV overproduction complement the growth defect of *topA* null mutants by relaxing negative supercoiling. In fact, topo IV plays a role in the regulation of global negative supercoiling (28).

Topo I relaxes negative supercoiling generated behind moving RNAPs during transcription (8). The failure to relax transcription-induced negative supercoiling in *topA* null mutants leads to hypernegative supercoiling and the formation of R-loops that can inhibit gene expression and activate DnaA-independent unregulated replication (29–35). In fact, overproduction of RNase HI, an enzyme degrading the RNA portion of a DNA:RNA hybrid in the R-loop, can compensate for the growth defect of *topA* null mutants (36). Moreover, the supercoiling activity of gyrase acting in front of replication and transcription complexes promotes R-loop formation both *in vitro* and *in vivo* (36,37). Thus, one major function of topo I in *E. coli* is the relaxation of transcription-induced negative supercoiling to inhibit R-loop formation.

Unlike *topA* null mutants, *topB* null cells grow as well as wild-type cells, but show a delayed and disorganized nucleoid segregation phenotype, as compared to wild-type cells (16). Considering that topo III is found associated with replication forks *in vivo* where its substrate, ssDNA, is present and that *in vitro* it has a strong decatenation activity that can fully support replication, it is likely that topo III plays a role in the removal of pre-catenanes. However, at least under normal growth conditions and when topo I is present, this role appears to be minor as compared to topo IV, the absence of which inhibits growth and causes severe chromosome segregation defects (the *par* phenotype (21)).

Initially, *topA topB* null mutants were shown to generate a RecA-dependent chromosome segregation and extreme cellular filamentation phenotype and were reported to be non-viable (38). Nevertheless, it was later found upon prolonged incubation that *topA topB* null transductants could be obtained (39). More recently, a correlation between R-loop-triggered RecA-dependent amplification in the chromosomal Ter region, cellular filamentation, chromosome segregation defects and growth inhibition phenotypes was demonstrated in *topA topB* null mutants (22). RNase HI overproduction was shown to correct these phenotypes and R-loop formation was detected in a *topA* null as well as in a *topA topB* null mutant, but at a higher level in the latter. Importantly, chromosomal DNA amplifications carrying *parC* and *parE* genes for topo IV were detected in every *topA topB* null mutant studied, irrespective of the *topA* or *topB* null alleles, and whether RNase HI was overproduced ((22) and see this work). In parallel to this study, a similar observation was reported for *B. subtilis* i.e., amplification of a DNA region carrying the *parEC* operon (for topo IV) allowing *topA topB* null mutants to survive (40).

Here, to better understand the relative contribution of the relaxase and decatenase activity in the essential function(s) of type 1A topos in bacteria, we sought to characterize the effect of deleting *topB* on supercoiling in *topA* null mutants and the mechanism by which topo IV overproduction allows *topA topB* null cells to survive. Our main findings are: 1-there is a substantial effect of the absence of topo III on hypernegative supercoiling in *topA* null mutants. This illustrates the significant backup roles that topo I and topo III can play for each other when one of them is absent. 2-the inhibition of R-loop formation requires a topo that can act locally (topo I or topo III, but not topo IV) to relax transcription-induced negative supercoiling. 3-topo IV overproduction is mostly required to solve R-loop- and NER-dependent topological problems related to replication in the absence of type IA topos. 4-excess R-loop formation is well tolerated in cells that have enough topoisomerase activity to support the ensuing replication-related topological stress.

## MATERIALS AND METHODS

### Bacterial strains and plasmids

The list of bacterial strains used in this study as well as the details on their constructions can be found in Table S1. This table also give the list of plasmids used in this work. Transductions with phage P1*vir* were done as described previously (41). PCR with appropriate oligonucleotides were performed to confirm the transfer of the expected alleles in the selected transductants. Strains JB136, JB335 and JB336 also carry a Δ*yncE* mutation that was introduced in the *gyrB*(Ts) strain before the Δ*topB* and *topA20*::Tn*10* alleles. This mutation in the Ter DNA region was introduced in the frame of our study aiming at the identification of loci that could affect Ter DNA amplification. It was found that this mutation had no effects on *topA topB* null cells growth, cell morphology or MFA profiles including Ter DNA amplification (Fig. S1, compare the MFA profile of JB137, *topA topB* and JB335, *topA topB* Δ*yncE*). The inversion in JB136 involves recombination between short homologous sequences (FRT, FLP recognition target) in the *yncE* and the *topB* loci that are the result of FLP recombinase activity to remove the Km^R^ cassette (42), that was introduced to substitute for these two genes during the process of JB121 strain construction. The inversion in JB136 was therefore named *IN(1.52-1.84)*. The mutations *uvrB24, uvrBC25* and *gyrA21* all appeared spontaneously in *topA topB* null mutants, as we confirmed by sequencing that they were not present in the *topB* null strains in which the *topA* null allele was introduced.

### Bacterial cells growth

Cells were grown overnight at 37°C on LB plates added with the appropriate antibiotics and supplements. Cell suspension were prepared, and aliquots were used to obtain an OD_600_ of 0.01 in fresh LB media. The cells were grown at the specified temperature to log phase at the indicated OD_600_.

### Plasmid extraction for supercoiling analysis

pACYC184Δ*tet*5’ DNA extraction for supercoiling analysis was performed by using the Monarch®Plasmid Miniprep Kit (NEB). Cells were grown at the indicated temperature to an OD_600_ of 0.4. We found that after the first centrifugation step, keeping the cell pellet on ice for 30 minutes instead of resuspending it immediately in the plasmid resuspension buffer did not affect the results. Chloroquine gel electrophoresis was done as previously reported (43). The gels were stained with SYBR™ Gold Nucleic Acid Gel Stain (ThermoFisher Scientific). The one-dimensional gels (7.5 μg/ml chloroquine) were photographed under UV light by using the Alphaimager HP. The two-dimensional gels (7.5 and 30 μg/ml chloroquine respectively in the first and second dimension) were photographed by using the Blue light transilluminator (ThermoFisher scientific).

### Spot tests

Cells were grown at 30°C to an OD_600_ of 0.6. Five μl of 10-fold serial dilutions were spotted on LB plates that were then incubated at 30°C.

### Detection of DnaA-independent replication

For the detection of DnaA-independent replication, EdU (ethynyl deoxyuridine) incorporation and click-labeling using the “Click-It®Alexa Fluor 448®Imaging kit” (Life Technologies, Molecular Probes) were done as previously described (34,44). Briefly, the cells were grown in LB medium at 30°C to an OD_600_ of 0.3. An aliquot of cells was used for EdU incorporation for 60 min to detect ongoing replication in log phase cells. To detect DnaA-independent replication the log phase cells were first treated with spectinomycin for two hours, to allow the termination of replication rounds initiated from *oriC*, before EdU incorporation for 60 min. After click-labeling, replication (EdU incorporation) was detected by flow cytometry using a FACS Canto II apparatus (BD), and the data were analyzed and generated using the FACS Diva software.

### Marker frequency analysis (MFA) by next generation sequencing (NGS)

MFA by NGS was performed essentially as described previously (22). Cells were grown at 30°C to an OD_600_ of 0.4. An aliquot of 10 ml of the cell culture was transferred in a tube filled with ice for genomic DNA extraction. Spectinomycin (400μg/ml) was added to the remaining cell culture and the cells were incubated for an additional two hours at the same temperature and an aliquot of 10 ml of the cell culture was taken and treated as above. Cells were recovered by centrifugation. Genomic DNA was extracted by using the GenElute bacterial genomic DNA kit (Sigma Aldrich) as described by the manufacturer except that the treatment to proteinase K was for 2 hours instead of 30 min. This modification eliminates artefacts in the MFA profiles around the *rrn* operons (45). Sequencing was performed by using Illumina NovaSeq 6000 (Génome Québec, Montréal, Canada) to determine sequence copy number. Bioinformatics analysis was performed at the Canadian Centre for Computational Genomic (C3G, McGill University, Montréal, Canada). The *E. coli* K12 W3110 genomic sequence AP009048.1 was used as the reference for the read mapping (15 to 20 million sequencing reads per sample). To reduce miscalculation of depth of coverage due to reads mapping at multiple places in the genome, a minimum mapping quality of 10 (phred scale based) was used for a read to be kept during the calculation of depth of sequencing. The number of reads were normalized against a stationary phase wild-type control to consider differences in read depth across the genome of cells. Enrichment in 500 bp windows (on average) across the genome (10,000 points) was calculated and loess regression curves were generated with loess_span parameters set to 0.1. To evaluate the level of additional replication in the chromosomal Ter region after the spectinomycin treatment, we first determined the difference between the highest loess value in the Ter peak and the lowest one on the right side of the peak after and before the addition of spectinomycin. We then subtracted the number obtained for the MFA profile before spectinomycin to the one after the treatment.

### Variant calling

The available paired end reads were trimmed of Illumina adaptors using Trimmomatic (v 0.35). Variant calling against the AP009048.1 reference sequence was performed individually for each sample using the trimmed forward and reverse Illumina paired end reads using default settings from the snippy algorithm (v 4.6.0). The core comparison file was then generated using the multiple outputs of snippy (Seemann T (2015), snippy: fast bacterial variant calling from NGS reads https://github.com/tseemann/snippy).

### qPCR

The genomic DNA for qPCR was prepared as described above for NGS. The Maxima SYBR Green qPCR Master Mix (2X) (ThermoFisher Scientific) was used with a Rotor-Gene 6000 (Corbett) apparatus. For each experiment two tubes were prepared that contained respectively 50 and 100 ng of genomic DNA. Experiments were repeated at least twice for each set of primers. The ratios were determined by using the 2^-Δct^ formula and standard deviations were calculated from these values. The primers were designed by using the PrimerQuest tool (IDT). Forward and reverse primer sequences (5’-3’) were GAGTACCGGGCAGACCTATAA and AGCCTACTTCGCCACATTTC for *lepA*, and CTGGACTCACTGGATAACCTTC and TGCGCCGTGTGGTAAATA for *qseC*.

### Fluorescence microscopy

Cells were grown to an OD_600_ of 0.8, centrifuged and resuspended in 1X PBS. Cells were mixed with polyformaldehyde 4% and a 2 μl aliquot was spread on a coverslip previously treated with (3-aminopropyl)triethoxysilane. Two μl of a 1/1000 dilution of SYTO 40 (ThermoFisher Scientific) was applied on the dried samples and the coverslip was deposited on a slide. Microscopy was performed by using a Nikon Ti2 apparatus with a Plan Apochromat 1.45 NA oil immersion 100x phase contrast objective and CFP filter set for epifluorescence imaging. The camera was a Hamamatsu Orca Flash 4.0. Images were processed by using the ImageJ software.

### Dot blots with S9.6 antibodies

Genomic DNA for the dot blots with S9.6 antibodies was prepared as described above for NGS except that the amount of RNase A added was reduced by 20-fold. Treatment of the genomic DNA to RNase III or RNase HI, as well as dot blotting were performed as described previously (22).

## RESULTS

### The absence of *topB* exacerbates the transcription-induced negative supercoiling phenotype of *topA* null mutants

It has been recently shown that deleting *topB* in *topA* null mutants exacerbates phenotypes of *topA* null mutants, one of them being R-loop formation (22). Whether topo III acts as a backup for topo I in inhibiting R-loop formation by relaxing transcription-induced negative supercoiling, or via other mechanism(s) is unknown. To address this point, the plasmid pACYC184Δ*tet*5’ was used to monitor transcription-induced negative supercoiling in our strains. This plasmid carries a deletion of the 5’ portion of the *tetA* gene that inactivates *tetA* gene translation (29). This deletion inhibits hypernegative supercoiling in *topA* null mutants related to membrane anchorage of the transcription complex via the *tetA* gene product. Transcription-induced negative supercoiling of this plasmid leads to R-loop formation (29). For this work, a pair of strains similar to the one employed in our previous study (22) was used (JB206, *topA20*::Tn*10 gyrB*(ts) and JB137, Δ*topB topA20*::Tn*10 gyrB*(Ts)). As shown before, these strains show better growth at 37°C owing to the partial inactivation of GyrB, reducing the negative supercoiling activity of gyrase. Upon transferring the strains from 37 to 30°C, gyrase is rapidly re-activated which stimulates hypernegative supercoiling that leads to growth inhibition (29,36).

To detect hypernegatively supercoiled plasmid DNA, electrophoresis in agarose gels in the presence of chloroquine at 7.5 μg/ml was performed. Under these conditions the more negatively supercoiled topoisomers migrate more slowly except for the fastest migrating band indicated by an arrow in Fig. 1A, which represents hypernegatively supercoiled plasmid DNA. Fig. 1A shows that hypernegatively supercoiled pACYC184Δ*tet*5’ accumulated significantly at 37°C only when both *topA* and *topB* were inactivated (compare lane 1, JB206 and lane 3, JB137). Thirty minutes after a transfer from 37 to 30°C, hypernegatively supercoiled pACYC184Δ*tet*5’ accumulated in both strains, although it is evident that the hypernegatively supercoiled topoisomers were proportionally more abundant when both *topA* and *topB* were absent (compare lane 2, JB206 and lane 4, JB137). The use of two-dimensional agarose gel electrophoresis in the presence of chloroquine confirmed the presence of hypernegatively supercoiled pACYC184Δ*tet*5’ following the 37 to 30°C temperature downshift, and the higher proportion of these topoisomers in the double *topA topB* null mutant (Fig. 1B, compare JB206 with JB137). The effect of deleting *topB* on the accumulation of hypernegatively supercoiled DNA in a *topA* null strain is possibly underestimated, because of the amplification of *parC* and *parE* genes in the *topA topB* null mutant, but not in the *topA* null strain. Indeed, by measuring the *qseC*/*lepA* ratio in which *qseC* is located in between *parC* and *parE* in the amplified DNA region, and *lepA* is close to, but outside the amplified region, it is possible to estimate the extent of *parC* and *parE* amplification, and therefore the level of topo IV overproduction. This procedure has been used previously and validated by MFA by NGS (22). The *qseC*/*lepA* ratio is respectively 1.3 and 4.9 for the *topA* null and *topA topB* null strains (Fig. 1C), suggesting that the level of topo IV protein production is higher by 3 to 4-fold in the *topA topB* null strain.

**Figure 1.**
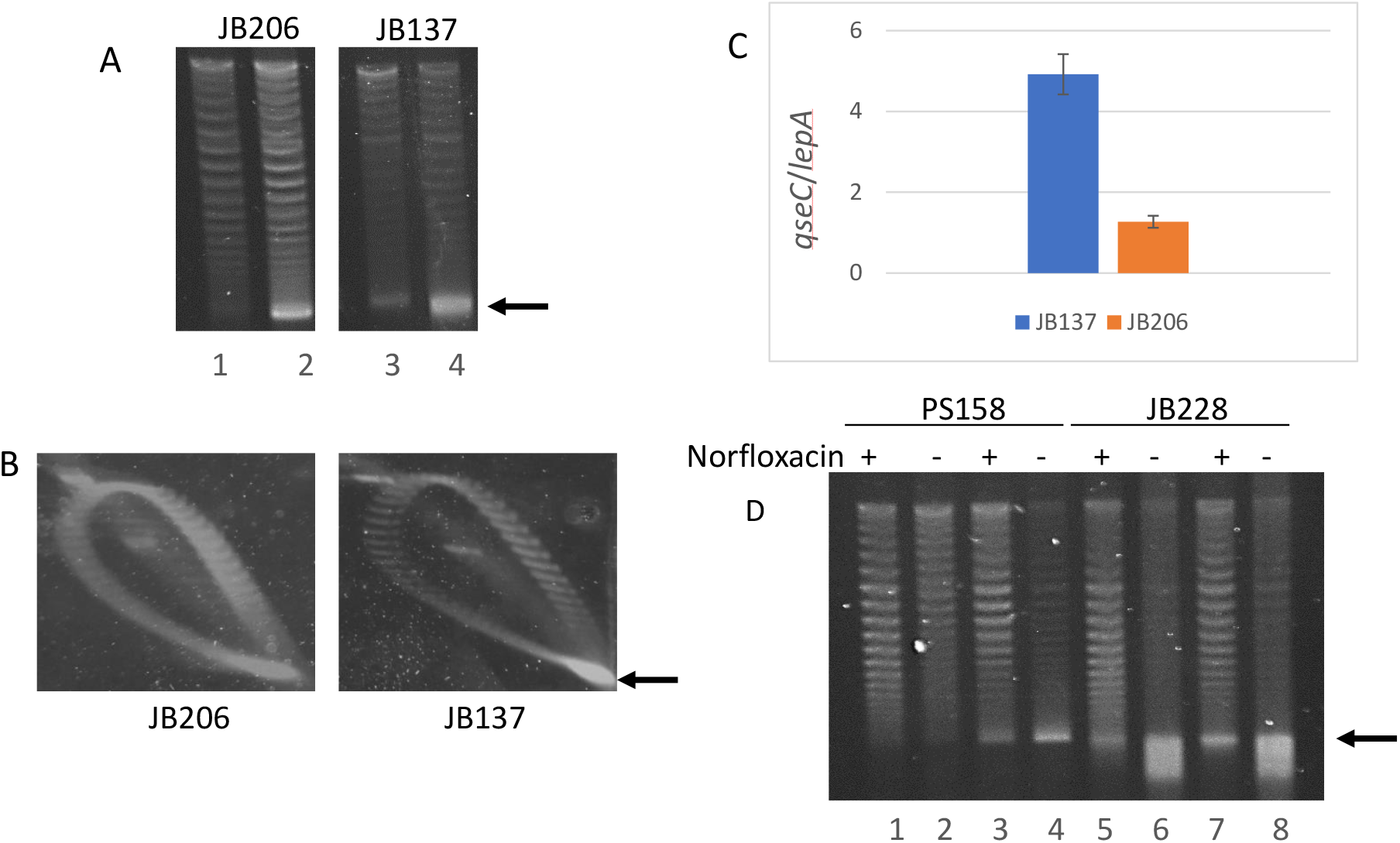
Hypernegative supercoiling is significantly stimulated by deleting *topB* in a *topA* null mutant. (A) One-dimensional chloroquine gel electrophoresis of pACYC184Δ*tet5’* extracted from JB206 (*topA20*::Tn*10 gyrB*(Ts))/pACYC184Δ*tet5’* and JB137 (*ΔtopB topA20::Tn10 gyrB*(Ts))/pACYC184Δ*tet5*’cells grown at 37°C to an OD_600_ of 0.4 (lanes 1 and 3), or 30 min after a transfer from 37 to 30°C (lanes 2 and 4). Aliquots of DNA samples used for lanes 2 and 4 in (A) were used for two-dimensional chloroquine gel electrophoresis in (B). (C) *qseC*/*lepA* ratio determined by qPCR. (D) One-dimensional chloroquine gel electrophoresis of pACYC184Δ*tet5’*. Cells of PS158 (*ΔtopA cys gyrA^L3^zei-723::Tn10 gyrB*(Ts))/pACYC184Δtet5’ and JB228 (PS158/pACYC184Δ*tet5ΔtopB*) strains were grown at 37°C to an OD_600_ of 0.4 and norfloxacin (60 μM final) was added or not as indicated. Aliquots of cells were taken for DNA extraction 30 min later (lanes 1, 2, 5 and 6) and the remaining cells were transferred to 30°C. 30 min later aliquots of cells were taken for DNA extraction (lanes 3, 4, 7 and 8). The strong signal at the bottom of lanes 6 and 8 correspond to hyper-negatively supercoiled DNA as it was fully relaxed by *E. coli* topo I in vitro (not shown). Arrows indicate hyper-negatively supercoiled DNA.

To better evaluate the real effect of topo III on hypernegative supercoiling, we used a pair of isogenic *topA* and *topA topB* null *gyrB*(Ts) strains that also carry the *gyrA^L83^* allele, which produces a gyrase resistant to high levels of norfloxacin (43). Since norfloxacin also inhibits topo IV, adding this drug to our isogenic strains will reveal the true effect of the absence of topo III on hypernegative supercoiling. Fig. 1D clearly demonstrates that topo IV inhibition had a much stronger effect in the double *topA topB* null mutant as compared to the *topA* null one (compare lanes 2 and 4, PS158 with lane 6 and 8, JB228, at 37°C and 37 to 30°C, + norfloxacin). This result reveals the dramatic effect of deleting *topB*, particularly at 37°C, on the accumulation of hypernegative supercoiling in a *topA* null strain. It also reveals that topo IV, when overproduced, can partly relax hypernegative supercoiling. In Fig. S2 we show that deleting *topB* had no effect on supercoiling in the *gyrB*(Ts) strain. Thus, topo III plays a backup role for topo I in the relaxation of transcription-induced negative supercoiling when topo I is absent, which likely explains the higher level of R-loop formation when both type 1A enzymes are lacking ((22) and see below).

### Human topo IB overproduction cannot substitute for topo IV overproduction in allowing *topA topB* null mutants to survive

Because topo IV did not appear to be very efficient in relaxing transcription-induced negative supercoiling, we considered the possibility that it is mostly required as a decatenase instead of a relaxase in allowing *topA topB* null mutants to survive. To test this hypothesis, we performed a simple genetic experiment to evaluate the ability of an efficient relaxase, but a poor decatenase, to substitute for topo IV in complementing *topA topB* null mutants. The spot test in Fig. 2A is a control showing that overproducing human topo IB (plasmid pET1B, JB244), an efficient enzyme to relax transcription-induced supercoiling but that plays no role in decatenation (2,46), can complement the growth defect of the single *topA* null mutant JB206 as well as topo IV overproduction (plasmid pET11-*parEC*, JB232), or even slightly better. This result demonstrates that pET1B produces enough topo IB to substitute for *E. coli* topo I in relaxing the growth-inhibitory excess negative supercoiling.

**Figure 2.**
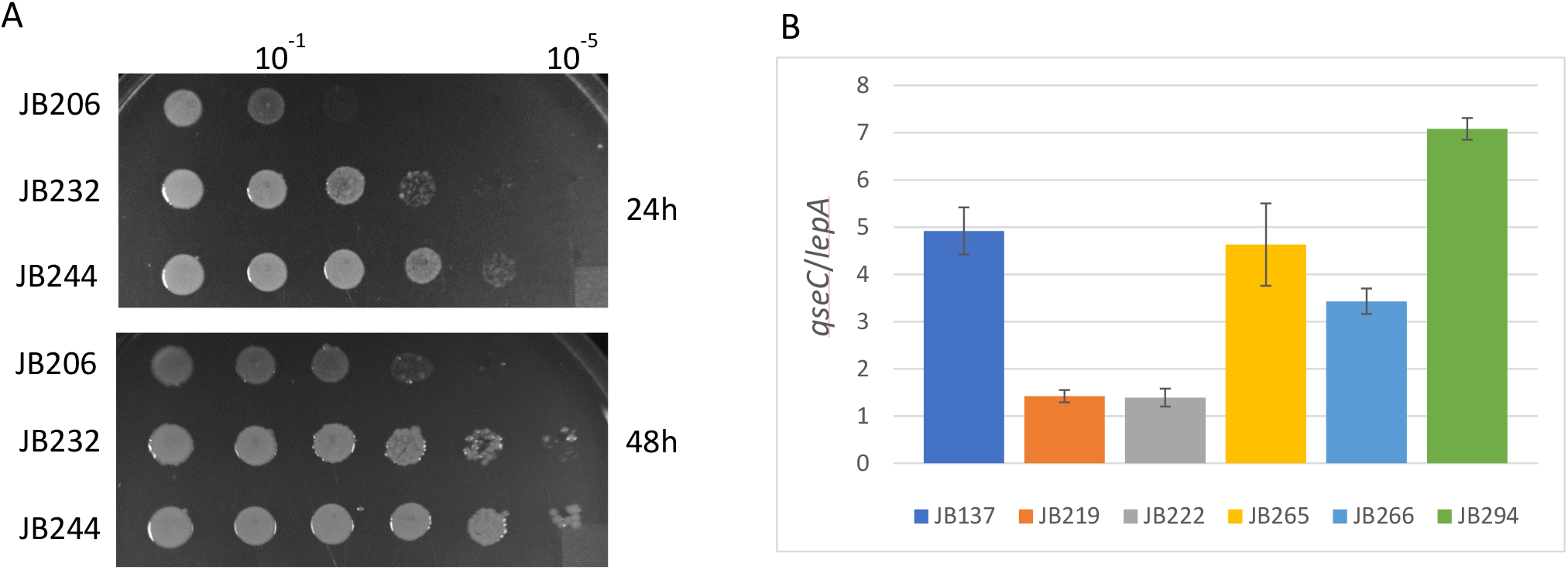
Human topo IB overproduction can substitute for topo IV overproduction in complementing the growth defect of *topA* null but not *topA topB* null mutants. (A) Ten-fold serial dilutions of log phase cell cultures of JB206 (*topA20*::Tn*10 gyrB*(Ts)), JB232 (JB206/pET11-*parEC*) and JB244 (JB206/pET1B) were spotted on an LB plate. Pictures were taken after 24 and 48 hrs of incubation at 30°C, as indicated. (B) *qseC*/*lepA* ratio determined by qPCR of genomic DNA from strains JB137 (*ΔtopB topA20::Tn10 gyrB*(Ts)), JB219 and JB222 (*ΔtopB topA20::Tn10 gyrB*(Ts)/pET11-*parEC*), and JB265, JB266 and JB294 (*ΔtopB topA20::Tn10 gyrB*(Ts)/pET1B).

Next, we performed a transduction experiment to introduce the *topA20*::Tn*10* allele in a *ΔtopB gyrB*(Ts) strain carrying either, pET11-*parEC*, pET1B or no plasmid. As expected, a selected plasmid-free *topA20*::Tn*10* Δ*topB gyrB*(Ts) transductant was found to carry an amplification of the *parC parE* DNA region, as shown by its *qseC*/*lepA* ratio of close to 5 (Fig. 2B, JB137). We tested two *topA20*::Tn*10* Δ*topB gyrB*(Ts) transductants carrying pET11-*parEC* and found that they did not have the *parC parE* amplification (Fig. 2B, JB219 and JB222, *qseC*/*lepA* ratio slightly above 1), as expected because topo IV can be overproduced from this plasmid. However, the three tested *topA20*::Tn*10* Δ*topB gyrB*(Ts) transductants with pET1B allowing topo IB overproduction, were found to carry a *parC parE* amplification, the level of which was higher if the transductant was selected at 30, instead of 37°C, a temperature that is more stringent for *topA20*::Tn*10* Δ*topB gyrB*(Ts) cells (Fig. 2B, JB265 and JB266, transductants obtained at 37°C, *qseC*/*lepA* ratios of 4.6 and 3.4 respectively, and JB294, a transductant obtained at 30°C, *qseC*/*lepA* ratio of 7.0). Thus, these results demonstrate that, as opposed to single *topA* null mutants, topo IB cannot substitute for topo IV overproduction to allow *topA topB* null mutants to survive. This result supports the hypothesis that topo IV overproduction complements an essential decatenase activity that is lacking due to the absence of topo I and III.

The *topA topB* null *gyrB*(Ts) transductant used in this study (JB137) and in the previous one (22) were all obtained at 37°C, a more permissive temperature than 30°C for their growth due to the presence of the *gyrB*(Ts) allele. Therefore, the level of amplification of the *parC parE* DNA region is probably sufficient for good growth at 37 but not at 30°C, where the growth was shown to be strongly affected and the extreme filamentation and nucleoid shape defects were fully expressed ((22) and see below). Here, we tested if further increasing the level of topo IV by introducing the pET11-*parEC* plasmid in JB137 strain (strains JB208) or overproducing topo IB from pET1B (strain JB252) could suppress these phenotypes. It was found that both the growth and cell morphology defects were suppressed by pET11-*parEC* and pET1B at 30°C, with pET1B being slightly better for the growth. Indeed, Fig. 3A shows that pET1B (JB252) was almost as good as pSK760 (RNase HI overproduction, JB393) to complement the growth defect of JB137 at 30°C, whereas pET11-*parEC* (JB208) was not as good but clearly better than JB137 without plasmid. Regarding cell morphology, Fig. 3B clearly show that pET11-*parEC* was as good as pET1B to correct the cellular filamentation phenotype of JB137. The effect of pET11-*parEC* and pET1B on pACYC184Δ*tet*5’ hypernegative supercoiling was also tested. Fig. 3C shows that both plasmids reduced hypernegative supercoiling, with pET1B being slightly better than pET11-*parEC*. These differences are better seen by looking at the relative proportion of the topoisomers in the different lanes, especially at 37°C. Thus, overproduction of topo IV at the appropriate level can efficiently complement phenotypes of *topA topB* null mutants. Topo IB overproduction is also very effective in complementing these phenotypes providing that the level of topo IV is high enough for *topA topB* null cells to survive (as in JB137).

**Figure 3.**
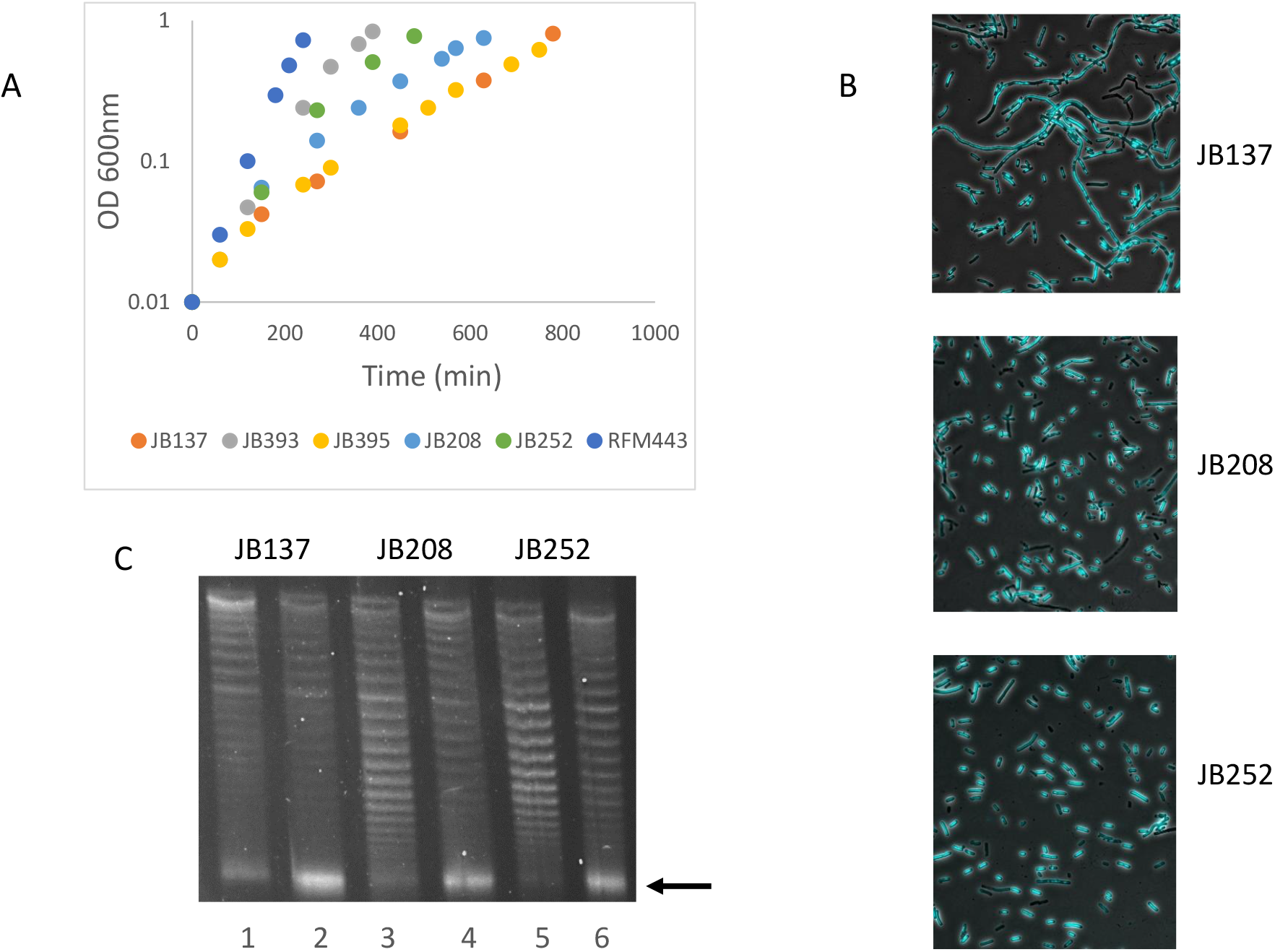
Effect of Human topo IB or topo IV overproduction from plasmids on the growth, supercoiling and cell morphology of the *topA topB* null mutant JB137. (A) Growth curves of RFM443 (wild-type), JB137 (*ΔtopB topA20::Tn10 gyrB*(Ts)), JB393 (JB137/pSK760), JB395 (JB137/pSK762c), JB208 (JB137/pET11-*parEC*) and JB252 (JB137/pET1B) strains at 30°C in LB. (B) Representative merged images of phase contrast and fluorescence pictures of SYTO-40-stained JB137, JB208 and JB252 cells grown at 30°C. (C) One-dimensional chloroquine gel electrophoresis of pACYC184Δ*tet5’* extracted from JB137/pACYC184Δ*tet5’*, JB208/pACYC184Δ*tet5’* and JB252/pACYC184Δ*tet5’* cells grown at 37°C to an OD_600_ of 0.4 (lanes 1, 3 and 5), or 30 min after a transfer from 37 to 30°C (lanes 2, 4 and 6). Arrows indicate hyper-negatively supercoiled DNA.

### Unlike topo IB overproduction, topo IV overproduction is unable to inhibit R-loop formation and RNase HI-sensitive replication

By using S9.6 antibodies recognizing DNA:RNA hybrids in dot-blot experiments, we have recently demonstrated the accumulation of R-loops in *topA topB* null cells (22). The dot-blot experiment in Fig. S3 shows, as expected, the accumulation of R-loops in the *topA* null (JB206) and *topA topB* null (JB137) strains used in the present study as well as in a *rnhA* null strain (MM84), but not in the control wild-type strain (RFM443). Fig. 4A shows that whereas topo IB overproduction (pET1B) inhibited the accumulation of R-loops in *topA topB* null cells (strain JB252), overproducing topo IV (pET11-*parEC*, strain JB208) had no significant effect on their accumulation. This is consistent with topo IB inhibiting R-loop formation by relaxing transcription-induced negative supercoiling in eukaryotes (46). Thus, inhibition of R-loop formation likely requires a topoisomerase acting locally to relax the transcription-induced negative supercoiling. This is consistent with topo IV being able to relax hypernegatively supercoiled DNA related to R-loop formation coupled to gyrase activity, but not to prevent its formation (33,47).

**Figure 4.**
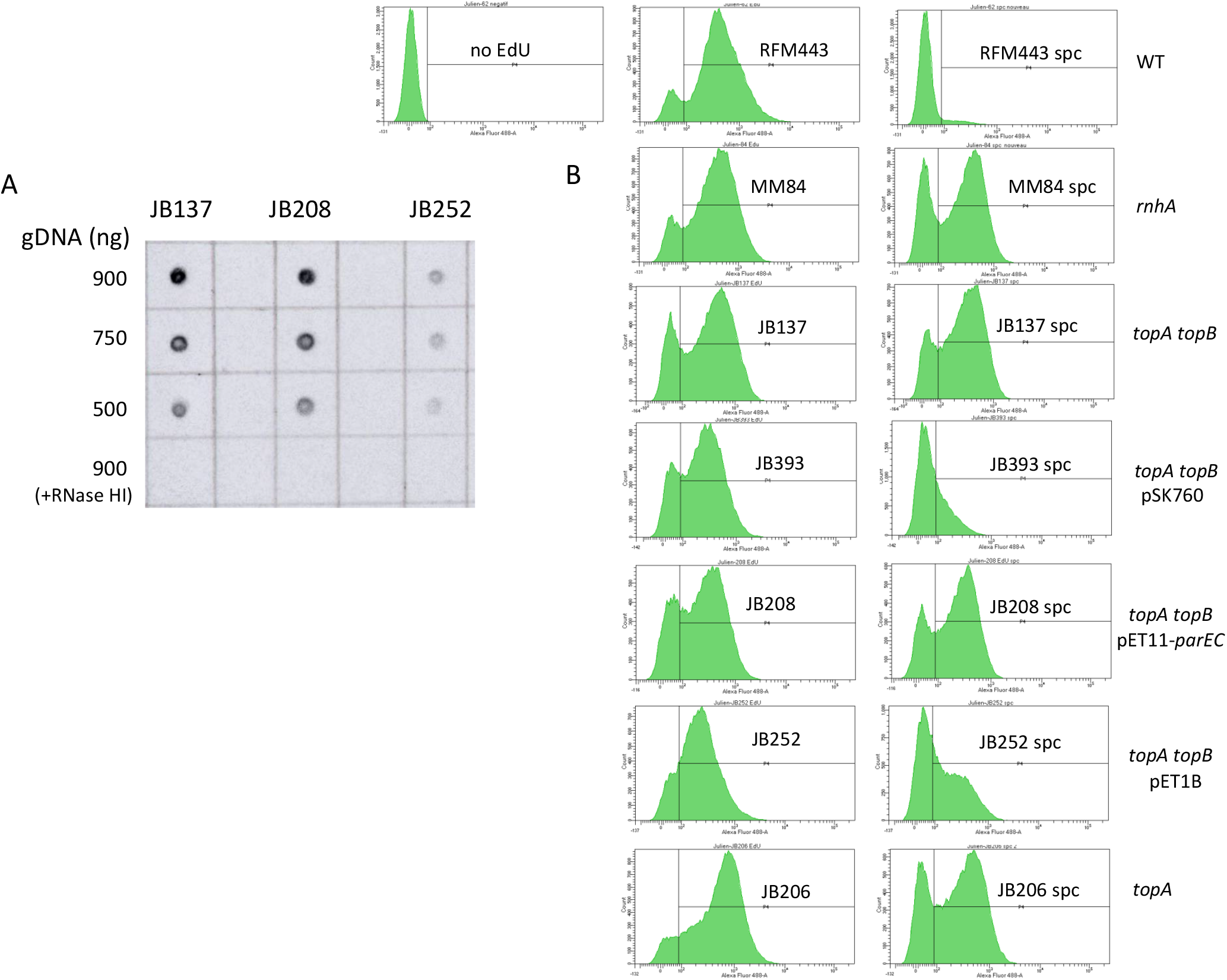
Overproduction of topo IV does not inhibit R-loop formation and RNase HI-sensitive replication. (A) Dot-blot with S9.6 antibodies of genomic DNA from JB137 (*ΔtopB topA20::Tn10 gyrB*(Ts)), JB208 (JB137/pET11-*parEC*) and JB252 (JB137/pET1B) cells grown at 30°C. The amount of genomic DNA spotted on the membrane is indicated. +RNase HI indicates that the genomic DNA was treated with RNase HI. (B) Flow cytometry showing DnaA-independent replication in RFM443 (wild-type), MM84 (*rnhA::cam*), JB137 (*ΔtopB topA20*::Tn*10 gyrB*(Ts)), JB393 (JB137/pSK760), JB208 (JB137/pET11-*parEC*), (JB137/pET1B) and JB206 (*topA20*::Tn*10 gyrB*(Ts)) cells grown at 30°C. no EdU is a control showing the position of the peak for cells that did not incorporated EdU (non-specific binding of the Alexa Fluor 448®dye). Spc, means that the cells were treated with spectinomycin for 2 hours before EdU incorporation.

Next, we analyzed the effect of topo IB or topo IV overproduction on DnaA-independent, RNase HI-sensitive replication in the *topA topB* null mutant, JB137. Initiation of DnaA-independent replication, also known as stable DNA replication (SDR), is resistant to protein synthesis inhibition as opposed to replication initiation via the normal DnaA/*oriC* system in bacteria. cSDR, for “constitutive stable DNA replication” was first described in *E. coli rnhA* (RNase HI) mutants and is therefore thought to originate from R-loops (48,49). We have recently developed a protocol to detect SDR that allowed the detection of such RNase HI-sensitive, DnaA-independent replication in both *topA* and *topA topB* null mutants ((34,44) and see Materials and Methods). In this protocol, log phase cells are first treated with spectinomycin for two hours to inhibit the initiation of replication from *oriC* and to allow the already initiated DnaA-dependent replication to be terminated. EdU, a thymidine analog, is then added to the cells and after one hour the cells are fixed, and the Alexa Fluor 488 dye is conjugated to EdU via the click chemistry. EdU incorporation is detected by flow cytometry.

Fig. 4B shows the results for the wild-type strain (RFM443). The left panel is the no EdU control where the peak represents non-specific binding of the dye to the cells. A similar peak was seen for all the strains when EdU was not added to the cells, and this control is therefore not shown for the other strains. The middle panel represents cells incorporating EdU (no spectinomycin, cells replicating their DNA). The right panel shows the cells treated with spectinomycin for 2 hrs before the addition of EdU. In the case of RFM443, the result demonstrates the absence of DnaA-independent replication in wild-type cells. DnaA-independent replication was readily detected in the *rnhA* null (MM84), *topA* null (JB206) and *topA topB* null (JB137) strains, with the highest level found in strain JB137 (Fig, 4B). This *dnaA-independent* replication is R-loop-dependent as it was efficiently inhibited by overproducing RNase HI (plasmid pSK760) in the *topA topB* null mutant (strain JB393). This SDR replication was previously shown to require RecA (48,49) and phenotypes of *topA topB* null cells, including DNA amplification in the Ter region, were efficiently corrected by deleting *recA* (22). Fig S4 shows that deleting *recA* inhibited the DnaA-independent replication in both *topA* (VU452, *topA recA* null) and *topA topB* (VU243, *topA topB recA* null) strains. Finally, in agreement with the results of the dot-blot experiments, overproducing topo IV from pET11-*parEC* (JB208) had no effect on DnaA-independent replication, whereas overproducing topo IB from pET1B (JB252) significantly reduced this replication (Fig. 4B). Thus, once again, these results support the hypothesis that topo IV does not complement *topA topB* null mutants via its relaxation activity, but instead via its decatenation activity.

### R-loop formation and RNase HI-sensitive DnaA-independent replication in *topB* null derivatives of widely used *topA* null mutants

So far, our data have been obtained with strains carrying a *gyrB*(Ts) mutation and in the W3110 background. W3110 is one of the two extensively used *E. coli* K12 strains, MG1655 being the other one. W3110 carries an inversion of a long segment of about 783 kb originating from *rrnD* and *rrnE* operons that includes *oriC* (50). As a result, this inversion modifies the positioning of the *oriC* region relative to the Ter region that may affect DNA amplification in the Ter region (51). In a previous study, *topA* null transductants were readily obtained in the MG1655 background and were though to be viable without compensatory mutations (52). We have obtained one of these transductants, strain VS111 that has been widely used for the study of transcription-induced negative supercoiling (53–55). The fact that this strain grew well led us to suspect that it has acquired the very frequent *parC parE* amplification. Indeed, the *qseC*/*lepA* ratio of VS111 is 3.8 (Fig. 5A), which demonstrates *parC parE* amplification. We introduced a *topB* null allele in VS111 and found that the strain still grew well, but a bit slower than VS111 (Fig. 5B, JB303, compare with VS111). This *topA topB* null strain (JB303) has a high *qseC*/*lepA* ratio of 6.9 (Fig. 5A), consistent with our finding that *topA topB* null cells need more topo IV than *topA* null cells to grow (22). Overproducing RNase HI from pSK760 (pSK762c is the control with a mutated and inactivated *rnhA* gene) had no effect on the growth of JB303 that already grew very well (Fig. 5B, compare JB303, with JB350 (JB303/pSK760) and JB352 (JB303/pSK762c)). This MG1655 *topA topB* null strain accumulated R-loops (Fig. 5C) and showed a high level of DnaA-independent RNase HI-sensitive replication (Fig. 5D, compare JB303 with JB350 and JB352). Thus, these results support those obtained with W3110 strains carrying a *gyrB*(Ts) allele (JB137) and clearly show that *topA topB* null strains overproducing topo IV to an optimal level, can grow well and tolerate a high level of R-loops and RNase HI-sensitive DnaA-independent replication. This is consistent with the hypothesis that topo IV overproduction provides the decatenation activity required to support this over-replication.

**Figure 5.**
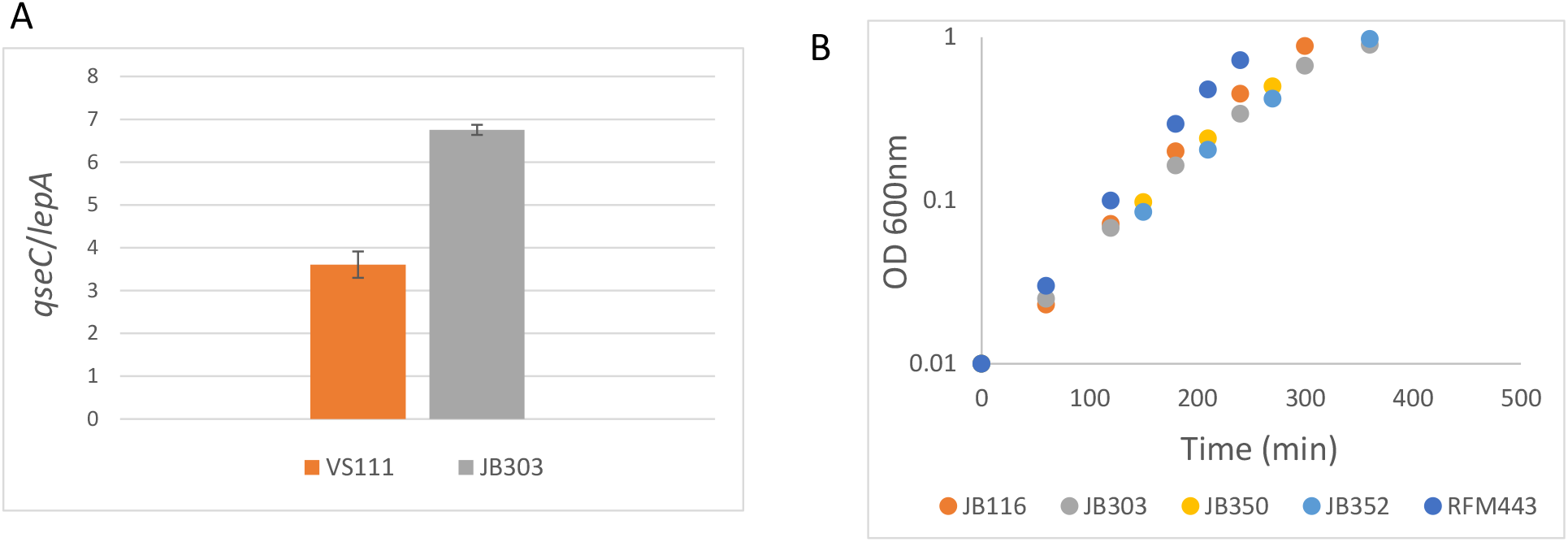
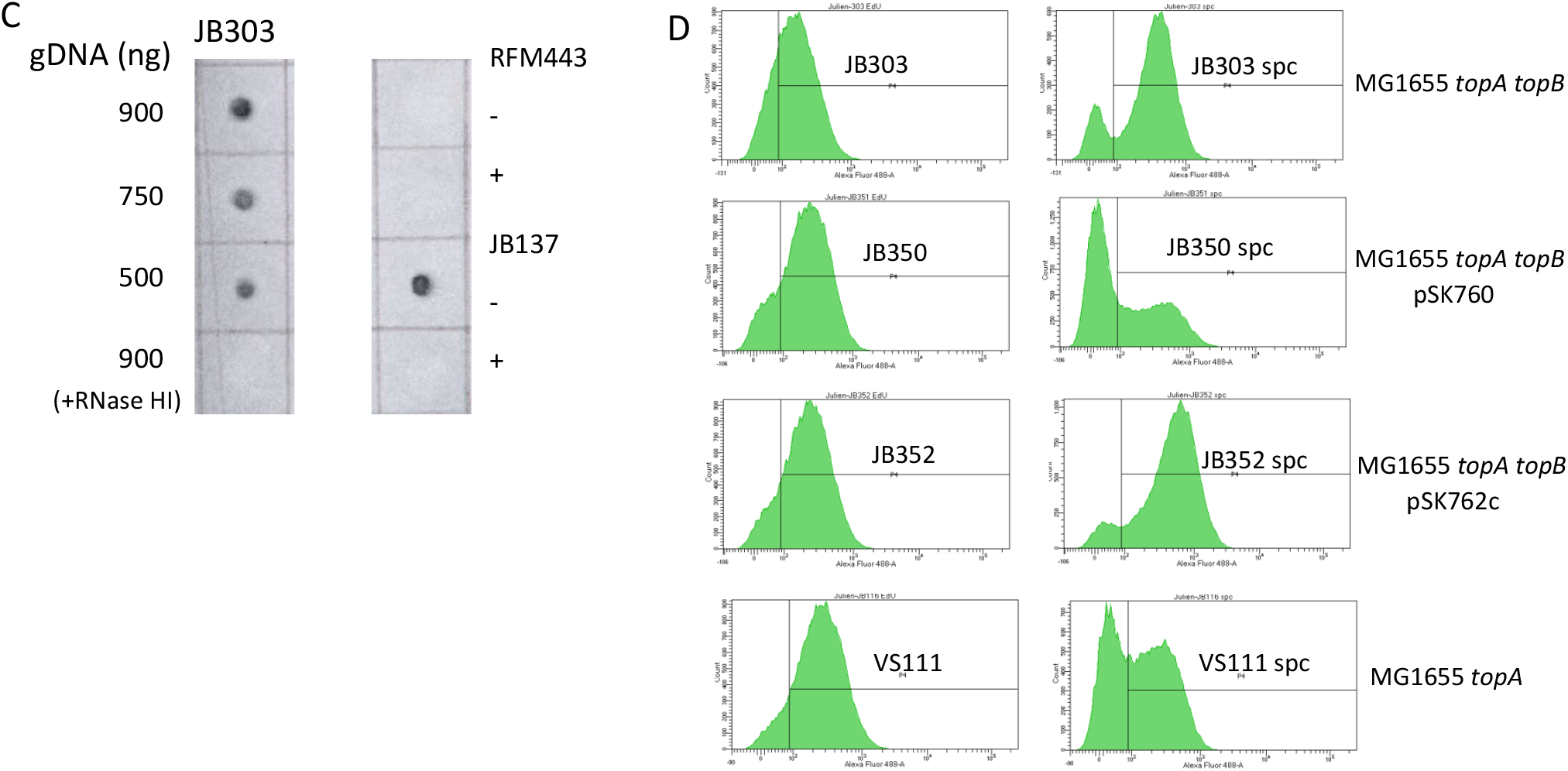
R-loop formation and RNase HI-sensitive DnaA-independent replication in a *topB* null derivative of MG1655 Δ*topA* null (strain VS111). (A) Growth curves of RFM443 (wild-type, from Fig. 3A), JB303 (MG1655 Δ*topA* ΔtopB), JB350 (JB303/pSK760) and JB352 (JB303/pSK762c) strains at 30°C in LB. (B) Dot-blot with S9.6 antibodies of genomic DNA from JB303 strain grown at 30°C. The amount of genomic DNA spotted on the membrane is indicated. +RNase HI indicates that the genomic DNA was treated with RNase HI. 900 ng of genomic DNA from RFM443 and JB137 (*ΔtopB topA20*::Tn*10 gyrB*(Ts)) were spotted on the membrane for comparison (− and + indicate that genomic DNA was respectively not treated or treated with RNase HI). (C) Flow cytometry showing DnaA-independent replication in JB303 (MG1655 Δ*topA* Δ*topB*), JB350 (JB303/pSK760), JB352 (JB303/pSK762c) and VS111 (MG1655 Δ*topA*) cells grown at 30°C. Spc, means that the cells were treated with spectinomycin for 2 hours before EdU incorporation.

We have also studied the effect of deleting *topB* on phenotypes of the widely used *topA* null strain DM800 carrying a naturally occurred compensatory *gyrB* mutation (*gyrB225*). This strain was used in the initial study showing that *topA* null mutants can survive owing to compensatory mutations reducing gyrase supercoiling activity (17,18). It was also used in the initial study of *topA topB* null mutants demonstrating that deleting *recA* corrected phenotypes of cells lacking type 1A topos (38). Here, we have constructed a *topB* null derivative of DM800 (JB305) and found that its growth could be slightly improved by overproducing RNase HI (Fig. S5A, compare JB305, JB354 (JB305/pSK760 and JB356 (JB305/pSK762c). JB305 grew slightly better than JB137 but more slowly than JB303 (see Fig. 3A, 5B and S5A). Deleting *topB* in DM800 strongly stimulated transcription-induced negative supercoiling and R-loop formation. Indeed, whereas no hypernegatively supercoiled pACYC184Δ*tet*5’ could be detected in DM800, such topoisomers were readily detected when *topB* was deleted (Fig. S5B, compare lanes 1 and 2, DM800 with lanes 3 and 4, JB305, 37°C and 37 to 30°C; note that the *gyrB* mutation of DM800 is not thermo-sensitive). Fig. S5C shows the significant effect of deleting *topB* on R-loop formation in DM800 (compare DM800 with JB305). DnaA-independent RNase HI-sensitive replication was also detected in DM800 and JB305, although at a lower level than in the other *topA topB* null strains JB137 and JB303, consistent with the presence of the *gyrB225* allele that is also expressed at 30°C (Fig. S5C, compare JB354 and JB356, and compare the low level of SDR replication in DM800, with the higher one in *topA* strains JB206 (Fig. 4B) and VS111 (Fig. 5D)). A *parC parE* amplification was not detected in strain JB305 (see below). Thus, a naturally occurring gyrase mutation presumably reduces enough DnaA-independent replication to prevent the critical overloading of the decatenation capacity of topo IV in the absence of type IA topos.

### MFA by NGS shows that both topo IB and topo IV overproduction reduces Ter region DNA amplification in *topA topB* null mutants

To better understand how topo IV overproduction could affect replication and complement the growth defect of cells lacking type 1A topos, we performed MFA by NGS as previously done to show RNase HI-sensitive amplification in the Ter-region of *topA topB* null mutants (22). The read count at every nucleotide position in the genome allows the high precision mapping of origins for bidirectional replication, the detection of DNA rearrangements such as duplications/amplifications and inversions and can reveal problems with the progression of replication forks. As done in our previous study, genomic DNA was extracted from log phase cells (OD_600_, 0.4) treated with spectinomycin (400 μg/ml) for 2 hrs. Genomic DNA was also extracted before the addition of spectinomycin to better reveal the actual replication taking place once the *oriC*-dependent one has, in principle, been completed (+ spectinomycin). The whole genome sequencing data were also used to perform variant calling to identify single nucleotide polymorphisms (SNPs) between our strains (Table S2). No SNPs were found between JB137, JB208 and JB252 strains. Importantly, apart from the *topA topB* null mutations, we were able to confirm that the *gyrB*(Ts) mutation was the only additional one present in these strains as compared to the isogenic wild-type strain RFM443.

By looking at the MFA profiles for strains JB137, JB208, JB252, JB303 and JB305 (Fig. 6), three observations can be made: 1-In the control with no added spectinomycin, the slope of the replication profiles from *oriC*, as expected, correlates with the strains’ growth rate as determined by the growth curves. 2-A *parC parE* amplification is detected in all the strains but JB305. 3-JB137 has a much higher Ter peak than the other strains that show better growth, either because they overproduce topo IV at a higher level (JB208 and JB303) or they reduce DnaA-independent (R-loop-dependent) replication (JB252 overproducing topo IB and JB305 that has a naturally occurred *gyrB* mutation). This chromosomal Ter region is normally the site of bidirectional replication forks fusion that allows replication to be completed. To ensure that replication termination take place in this region where the final chromosome decatenation by topo IV occurs, replication forks are trapped within the Ter region by Tus protein that binds to polar Ter sequences (10 of them, *TerA* to *TerJ*, (56)). Bidirectional replication taking place at the Ter region in *topA topB* null mutants shows that its left-(counterclockwise) and right-(clockwise) moving forks are respectively arrested at *TerA* and *TerB* (Fig. 6). Fig. 6 also reveals that the size of the amplified *parC parE* region is much shorter for the MG1655 background (JB303) as compared to the other strains in the W3110 background (JB137, JB208 and JB252). This may explain in part why the *topA topB* null mutant grows better in the MG1655 background.

**Figure 6.**
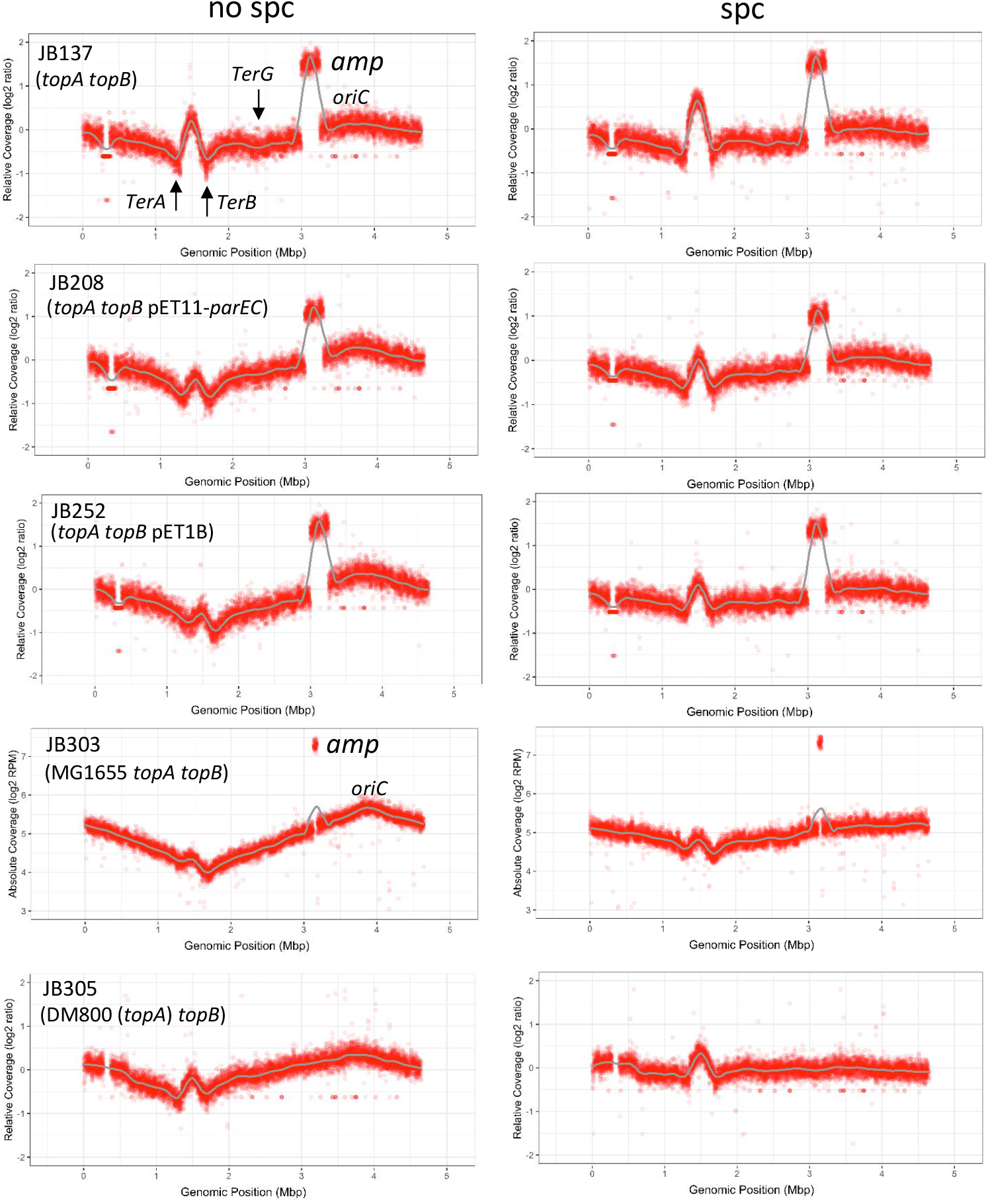
MFA by NGS shows that both topo IB and topo IV overproduction reduces Ter region DNA amplification in *topA topB* null mutants. MFA by NGS of genomic DNA extracted from JB137 (Δ*topB topA20*::Tn*10 gyrB*(Ts)), JB208 (JB137/pET11-*parEC*), JB252 (JB137/pET1B), JB303 (MG1655 Δ*topA* Δ*topB*) and JB305 (DM800 (Δ*topA*) Δ*topB*) cells grown at 30°C and treated (spc) or not treated (no spc) with spectinomycin for 2 hours. The absolute read counts (Log2) were plotted against chromosomal coordinates (W3310 genomic sequence AP009048.1). The gray line is the loess regression curve. *amp* indicates the amplified *parC parE* region. The gap at position around 0.3 in all the strains but JB303, corresponds to the Δ(*codB-lacI)3* deletion in these strains.

The level of additional replication in the Ter region after the spectinomycin treatment was evaluated for strains JB137 and JB208, by determining the increase in the Ter peak height two hours after spectinomycin addition (see Materials and Methods). The values for JB137 and JB208 are very similar, 0.234 and 0.210 respectively, despite the much lower Ter peak for JB208. This strongly suggested that the rate of DNA synthesis in the Ter region was very similar in the two strains and therefore indicated that topo IV overproduction at a high level (JB208) acted by reducing the amount of DNA accumulated in Ter region. Thus, topo IV and R-loop inhibition may act in the same pathway, the latter by preventing Ter amplification and the former by acting later to reduce Ter amplification.

### A *tus* deletion eliminates the Ter peak, improves growth, and reduces cellular filamentation of a *topA topB* null mutant

If the Ter peak is problematic in JB137, deleting the *tus* gene to inactivate Ter barriers including *TerA* and *TerB* should improve the growth and correct the filamentation phenotype of *topA topB* null cells. Fig. S6A shows that deleting *tus* completely eliminated the peak in the Ter region of the *topA topB* null strain. The complete absence of a Ter peak may suggest that the over-replication (amplification) in Ter does not directly involve the use of a conventional origin of replication, but instead an alternative mechanism, such as the collision of replication forks with Ter/Tus barriers that triggers re-replication in the reverse orientation, as previously suggested (51,57,58). Regardless of the mechanism at play, our results show that deleting *tus* (JB260) improved the growth (Fig. S6B) and considerably reduced cellular filamentation (Fig. S6C) of a *topA topB* null mutant. In fact, JB260 grew as well as JB137/pET11*parEC* (JB208, Fig. 3A and JB260, Fig. S6B). However, *parC parE* amplification was still required for the growth of the mutant as shown from the MFA profile (Fig. S6A, *amp*). These results shows that the requirement for a minimal level of topo IV overproduction for the survival of *topA topB* null mutants is not related to Ter DNA amplification but that at a higher level, topo IV overproduction significantly improves the growth of such mutants by correcting a decatenation problem in Ter DNA region (see Discussion).

### *topA topB* null mutants without *parC parE* amplification carry *uvrB* or *uvrC* mutations

To further understand the requirement for topo IV overproduction in decatenation for the survival of *topA topB* null cells, we searched for the presence of mutations other than a *parC parE* amplification, that could allow the survival of cells lacking type IA topos. We have previously shown the absence of a *parC parE* amplification in a *topA topB* null strain carrying a *dnaT* mutation (*dnaT18::aph*) that was initially isolated as a suppressor mutation of the growth defect and cellular filamentation phenotype of a *topA rnhA* null mutant (34,35,59). This mutation is an insertion of an *aph* cassette (kanamycin resistance) in the promoter region of the *dnaTC* operon that reduces by 5-fold the expression of *dnaT* but has no effect on *dnaC* expression (22). This *dnaT18::aph* mutation significantly reduces SDR in a *rnhA* null mutant (34). Moreover, unlike a *dnaT* null mutation that is barely viable, a wild-type strain carrying our *dnaT18::aph* mutation grows normally. DnaT is an essential component of the PriA-dependent primosome involved in replication restart, in cSDR (DnaA-independent replication from R-loops) and in the RecA-dependent RecBCD pathway of homologous recombination (34,48,60). It is also required for the DNA amplification observed in chromosomal Ter region of *recG* mutants (61).

Upon re-streaking on plates our original *topA topB dnaT18::aph* mutant (strain VU441, (35)), we noticed a mixture of small and large colonies. The colonies were purified to generate strain JB325 (small colonies) and JB326 (large colonies). The growth curve of Fig. 7A shows that indeed JB326 grew better than JB325 and that both strains grew better than JB137 despite the absence of a *parC parE* amplification in the *dnaT* mutants. Interestingly, we found that both JB325 and JB326 carried a *gyrA* mutation (Table S2; this mutation was named *gyrA21*) that had no effect on hypernegative supercoiling (Fig S7A, compare JB137 and JB325). Importantly, one additional mutation was found in JB326, in *uvrB* (Table S2 and see below; this mutation was named *uvrB24*), which is therefore likely responsible for its better growth relative to JB325. This mutation had no effect on R-loop formation as shown from dot-blot experiments with total genomic DNA (Fig. S7B, compare JB325 and JB326). MFA profiles of JB325 and JB326 demonstrated a significant reduction in the level of Ter DNA amplification and confirmed the absence of *parC parE* amplification (Fig. 7B). Furthermore, values of 1.42 and 1.22 were obtained by quantifying the increase in the peak height two hours after spectinomycin addition, respectively for JB325 and JB326 (see Materials and Methods). These values are significantly lower than those for JB137 and JB208 strains (2.34 and 2.10, see above). This is consistent with the prediction that the *dnaT18::aph* mutation reduces DNA synthesis in the Ter region. We also note that because this mutation reduced Ter DNA amplification, it might also have precluded the accumulation of a *parC parE* amplification and therefore added a selective pressure for the occurrence of the *gyrA21* mutation. This would suggest that a fully active primosome is required in *topA topB* null mutants in other circumstances (see Discussion).

**Figure 7.**
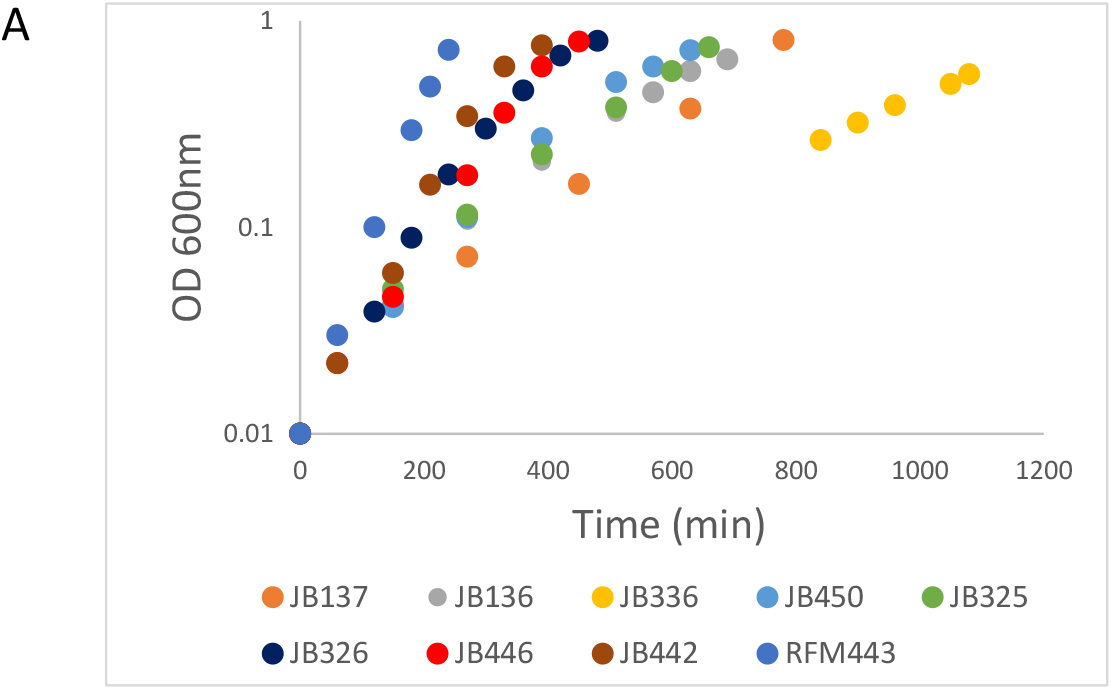
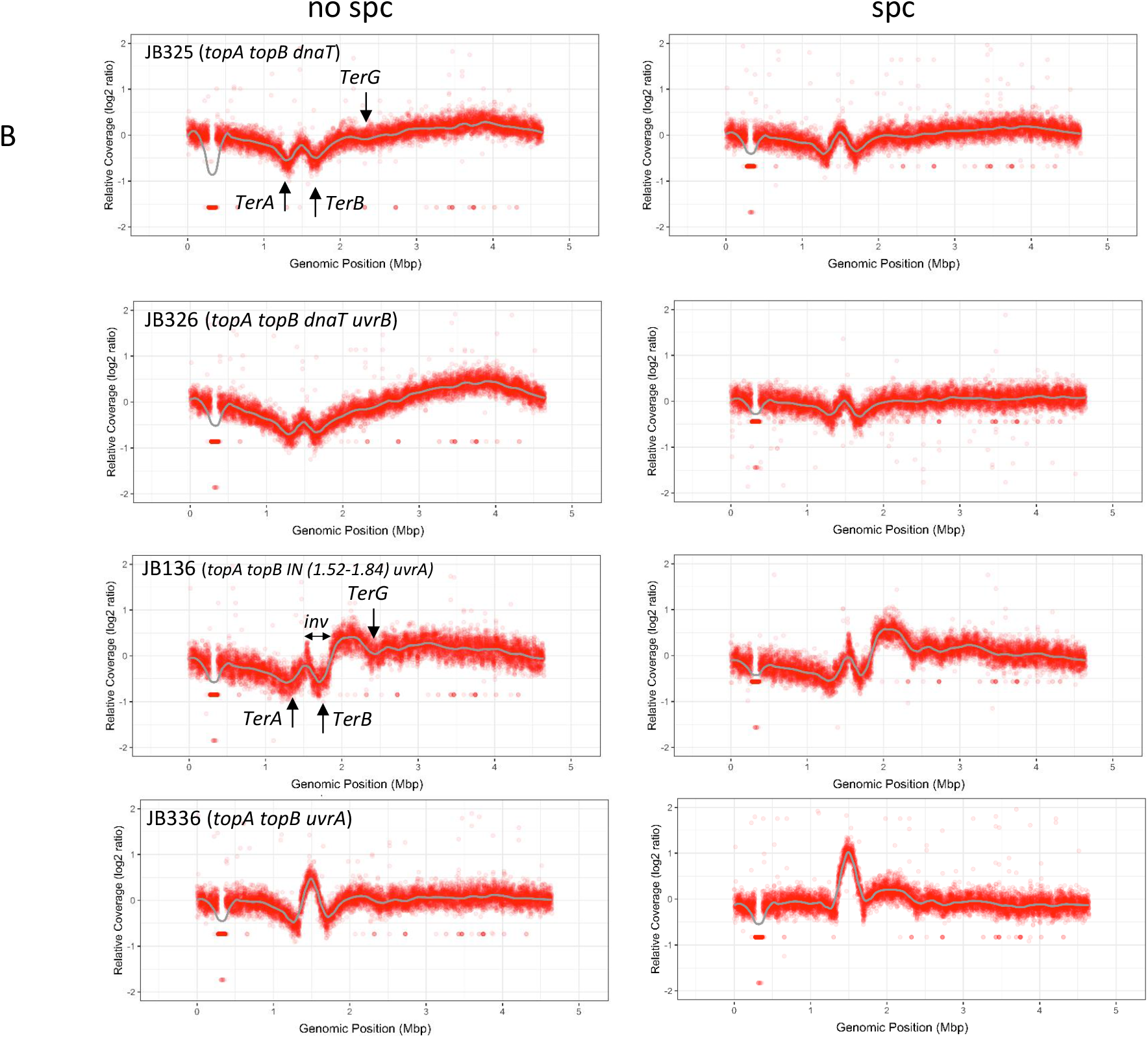
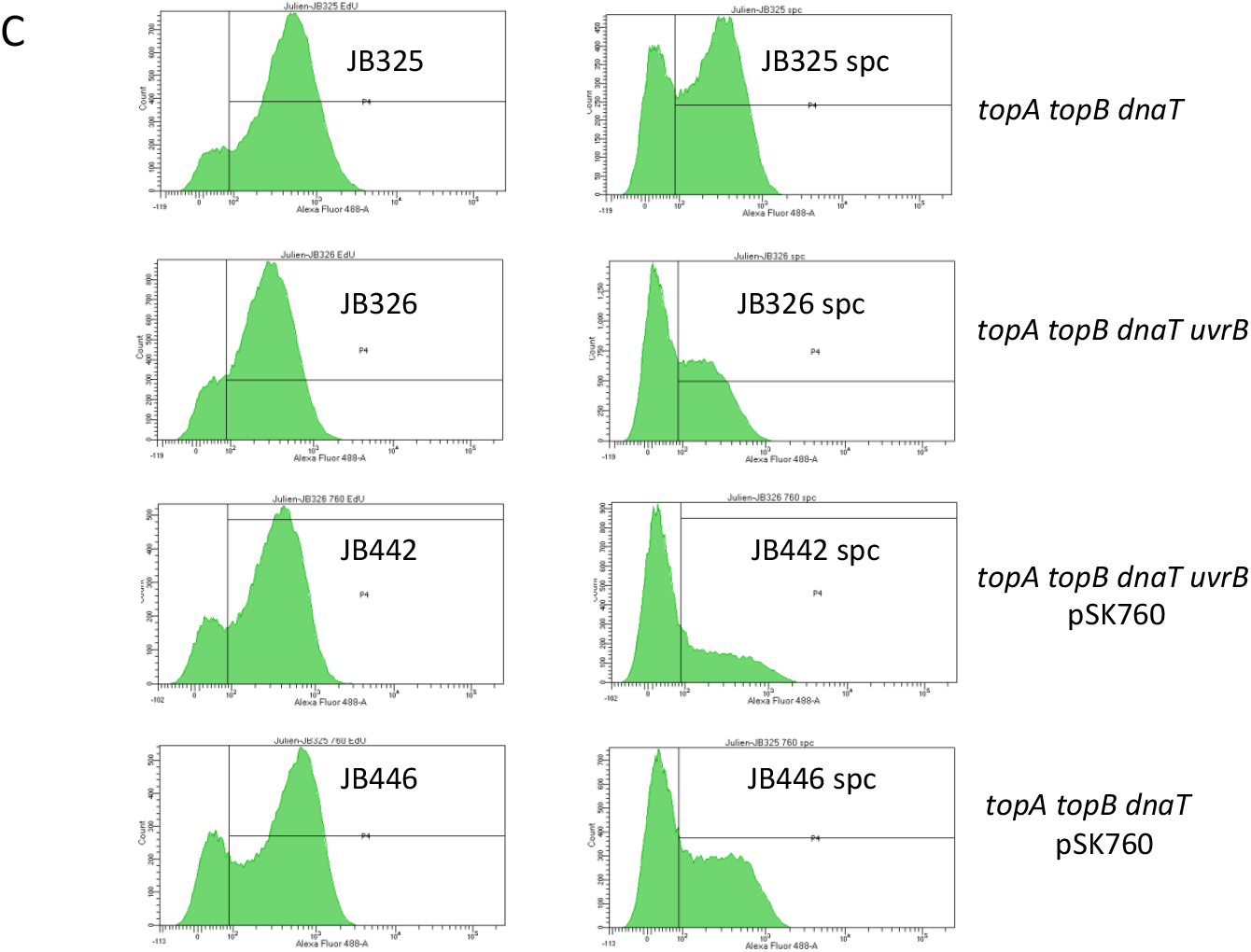
Effects of dnaT18::aph, uvrB24 and uvrC25 mutations on phenotypes of *topA topB* null cells not carrying *parC parE* amplifications. (A) Growth curves of RFM443 (wild-type, from Fig. 3A), JB136 (*ΔtopB topA20::Tn10 gyrB*(Ts) *IN(1.52-1.84) uvrC25*), JB137 (*ΔtopB topA20::Tn10 gyrB*(Ts)), JB325 (*ΔtopB topA20::Tn10 gyrB*(Ts) *gyrA21*), JB326 (*ΔtopB topA20::Tn10 gyrB*(Ts) *gyrA21 uvrB24*), JB336 (*ΔtopB topA20::Tn10 gyrB*(Ts) *uvrC25*), JB442 (JB326/pSK760), JB446 (JB325/pSK762c) and JB450 (JB336/pSK760) strains at 30°C in LB. (B) MFA by NGS of genomic DNA extracted from JB325 (*ΔtopB topA20::Tn10 gyrB*(Ts) *gyrA21*), JB326 (*ΔtopB topA20::Tn10 gyrB*(Ts) *gyrA21 uvrB24*), JB136 (*ΔtopB topA20::Tn10 gyrB*(Ts) *IN(1.52-1.84) uvrC25*) and JB336 (*ΔtopB topA20::Tn10 gyrB*(Ts) *uvrC25*), cells grown at 30°C and treated (spc) or not treated (no spc) with spectinomycin for 2 hours. See legend to Fig. 6 for more details. (C) Flow cytometry showing DnaA-independent replication in JB325 (*ΔtopB topA20::Tn10 gyrB*(Ts) *gyrA21*), JB326 (*ΔtopB topA20::Tn10 gyrB*(Ts) *gyrA21 uvrB24*), JB442 (JB326/pSK760) and JB446 (JB325/pSK762c) cells grown at 30°C. Spc, means that the cells were treated with spectinomycin for 2 hours before EdU incorporation.

UvrB is part of the bacterial nucleotide excision repair (NER) pathway that also includes UvrA and UvrC proteins (62,63). This system is extraordinary versatile as it recognizes and removes a wide variety of lesions that cause an important distortion of the double helix (64). The *uvrB* mutation in JB326 is a C to A substitution leading to a phenylalanine (conserved) to leucine change at position 89 of UvrB that is very close to the β-hairpin domain of UvrB (Y92 to K111). The β-hairpin separates the DNA strands at the site of DNA damage (65). It is required for DNA binding, damage processing and UvrC-mediated incisions (66).

By looking more closely at the + and – spectinomycin MFA profiles of JB325 and JB326 (Fig. 7B) we noticed that only strain JB326 gave a flat profile after spectinomycin treatment. This is normally expected if *oriC*-dependent replication is fully completed. Thus, the difference between the + and – spectinomycin profiles is much more important for JB326 which shows better growth and therefore shows more *oriC*-dependent replication in the absence of spectinomycin. This difference in the profiles of JB325 and JB326 is more evident by looking at Fig. S8 and it shows that replication fork progression is somehow problematic in JB325. In agreement with this hypothesis, we found a higher level of replication in JB325 as compared to JB326 two hours after spectinomycin treatment (Fig. 7C). This excess replication in JB325 may indeed correspond to ongoing replication from *oriC* that is difficult to complete. The remaining replication, also present in JB326, is highly sensitive to RNase HI overproduction and is therefore related to R-loops (Fig. 7C, compare JB326 and JB442 (JB326/pSK760)). As stated above, because the *dnaT18::aph* mutation only reduces the amount of DnaT protein, this remaining replication might be PriA-dependent from R-loops. RNase HI overproduction only slightly improved the growth of JB326 that was already very good (Fig. 7B, JB326 and JB442 (JB326/pSK760)). pSK760 somehow appeared to be toxic for JB325 as it has been difficult to obtain a JB325/pSK760 clone that could regrow. We analyzed one of them and found that RNase HI overproduction significantly stimulated the growth of JB325 (Fig. 7A, JB325 vs JB446 (JB325/pSK760). However, RNase HI overproduction only slightly reduced replication after the spectinomycin treatment, indicating that the NER-dependent replication fork impediment was still present (Fig. 7C, compare JB442 spc with JB446 spc). Thus, these results reveal a NER-dependent inhibition of replication fork progression in *topA topB* null cells. However, this phenotype is not present in *topA topB* null cells carrying a *parC parE* amplification, as in this case RNase HI overproduction almost fully eliminated the spectinomycin-resistant replication (Fig. 4B, JB393, JB137/pSK760).

In two independent transduction experiments involving the introduction of the *topA20*::Tn*10* allele in the *topB gyrB*(Ts) strain, we were able to obtain two *topA topB* null transductants that do not carry the *parC parE* amplification. Interestingly, these two transductants, strains JB136 and JB336 were found to carry the same *uvrC* mutation (see below; this mutation was named *uvrC25*). The peak in the Ter region of JB336 is as high as the one in JB137 (compare Fig. 7B, JB336 and Fig. 6, JB137). JB136 carries an inversion that changes the position of the amplification normally located in the Ter region (see Materials and Methods). JB136 grew a bit better than JB137, despite the absence of a *parC parE* amplification and grew much better than JB336 that barely grew (Fig. 7A). Furthermore, cells of JB136 do not form long filaments as is the case for JB137 (Fig. S9A). These results with JB136 further support the assumption that both the filamentation and growth defects of *topA topB* null mutants are largely related to DNA amplification in the Ter region. The inverted region in JB136 also includes *TerB*, so that this site now becomes the left boundary of the amplification whereas *TerG*, a weaker barrier (), is now the right boundary (Fig. 7B, JB136). The high level of DNA replication on the right side of *TerG* illustrates both the weak activity of this site as a barrier and the high level of replication originating from the *TerB TerG* interval. Strikingly, the peak height between *TerB* and *TerG* in JB136 is very similar to the peak height between *TerA* and *TerB* in JB137 (compare JB137, Fig. 6, with JB136, Fig. 7B), despite the lack of significant peaks in these areas in the absence of a *Ter*/Tus barrier (JB260, *topA topB tus*, no prominent peaks between *TerA* and *TerB* and JB137, *topA topB*, no prominent peaks in the area between *TerB* and *TerG*). This supports the hypothesis that amplification in both JB137 and JB136 does not directly involve the use of a conventional origin of replication, but instead an alternative mechanism (51,57,58).

The *uvrC25* mutation of JB136 and JB336, a T to A substitution (coding strand) at position 884, leads to a change from valine to glutamate at position 295 of UvrC (Table S2). The presence of an *uvr* mutation in these strains as well as in JB326, strongly suggests that it is related to the lack of a sufficient amount of topo IV to solve topological problems associated with NER-dependent replication fork impediments. Interestingly, we found that overproducing RNase HI (from pSK760) strongly stimulated the growth of JB336 (Fig. 7A, strain JB450 vs JB336). This in in contrast with the observation that, in the absence of *uvr* mutation, the few *topA20*::Tn*10* transductants of a *topB gyrB*(Ts) strain carrying pSK760 were shown to carry a *parC parE* amplification (Fig. S9B, compare JB511 (transductant with pSK760) with JB512 (transductant with pSK762c), JB136, JB336 and JB450 (JB336/pSK760)). This may suggest that RNase HI overproduction is somehow toxic when either the level of topo IV activity is too low or the NER system is fully active (see Discussion). We conclude that *uvr* mutations occur in the absence of *parC parE* amplification to allow *topA topB* null mutants to survive. Furthermore, our results suggest that the NER-dependent inhibition of replication fork progression somehow involves topological stress and makes RNase HI overproduction toxic.

## DISCUSSION

Because topos of the type IA family have unique substrates, are ubiquitous and were likely present very early in evolution, they were presumed to be essential enzymes. Therefore, as is the case for bacterial gyrase for example, it was assumed that their absence could not be compensated by any type of mutation including those modifying the activity of other topos. However, it was recently shown both in *B. subtilis* and in *E. coli* that their absence could be compensated by overproducing topo IV (22,40). These results implied that the activity of a topo acting on ssDNA is not absolutely required for cell growth. Since both topo IV and bacterial type IA topos can decatenate as well as relax DNA *in vitro* and *in vivo* it is not obvious to pinpoint the essential(s) activity(es) that topo IV complements in *topA topB* null cells. Here our data indicate that at low level of overproduction (e.g., strain JB137), topo IV allows *topA topB* null cells to survive by acting as a decatenase to prevent a toxic NER-dependent topological stress. At a higher level of overproduction (e.g., strains JB208 and JB303), topo IV stimulates the growth of *topA topB* null cells by solving decatenation problems related to Ter DNA amplification.

That topo IV does indeed act as a decatenase and not by relaxing negative supercoiling is mostly supported by these findings: 1-Overproduction of human topo IB, an enzyme that can efficiently relax but not decatenate DNA, can substitute for topo IV overproduction in complementing the growth defect of single *topA* null but not double *topA topB* null mutants. Furthermore, topo IB overproduction can significantly stimulate the growth of a *topA topB* null mutant, but only if topo IV is overproduced at a level allowing the double mutant to survive. 2-Unlike topos of the type IA and IB families that inhibit R-loop-dependent unregulated replication by relaxing transcription-induced negative supercoiling to prevent Ter DNA amplification and its associated chromosome segregation and growth defect phenotypes, topo IV overproduction acts at a later stage by directly solving the segregation defect. 3-The study of *topA topB* null mutants not carrying *parC parE* amplification reveals a NER-related defect in replication fork progression. These results also imply that type IA topos, by inhibiting R-loop formation and unregulated replication and by performing decatenation, respectively prevent and solve toxic topological problems. Furthermore, another important conclusion that can be reached from this work is that extensive R-loop formation and its associated DnaA-independent replication are well tolerated in cells that have enough topoisomerase activity to support the topological stress generated by this replication (e.g., strains JB208 and JB303).

### Ter DNA amplification and decatenation problems during replication termination in *topA topB* null mutants

Our MFA results show that the presence of a high peak in the chromosomal Ter region of *topA topB* null mutants correlates with poor growth, cellular filamentation and chromosome segregation defects. By performing MFA and qPCR, we have previously shown that a small peak was present in the Ter region of single *topA* null but not *topB* null mutants (22). This indicated that Ter amplification is triggered by the absence of topo I and dramatically increased by the additional absence of topo III. That *topA* deletion triggers the amplification is consistent with our data demonstrating that by relaxing transcription-induced supercoiling, topo I prevents R-loop formation and its associated DnaA-independent replication that is responsible for Ter DNA amplification. In this context, the significant increase in hypernegative supercoiling when *topB* is deleted in *topA* mutants could explain the high Ter peak in *topA topB* null mutants. However, we found that the level of both R-loop formation and R-loop-dependent replication in single *topA* null mutants is already very high and only slightly enhanced by deleting *topB* (R-loop formation, Fig. S3 and R-loop-dependent replication, Fig. 4B, JB206 vs JB137 and R-loop-dependent replication, Fig. 5D, VS111 vs JB303). Therefore, this backup role of topo III in relaxing transcription-induced negative supercoiling is not the main factor explaining the dramatic effect that deleting *topB* had on the Ter peak level in *topA* null mutants. In fact, our data are more compatible with the role of topo III in decatenation (7,16) that may also involve topo I as a backup when topo III is absent. In *topA topB* null mutants, this role would be compensated by topo IV overproduction at the appropriate level (JB208: JB137/pET11-*parEC* and JB303: MG1655 *topA topB*).

Unlike topo IB, gyrase mutations or RNase HI overproduction that prevent Ter DNA amplification, topo IV acts after Ter amplification to reduce it. Thus, the rate of Ter amplification is the same, whether topo IV is overproduced or not. Our results also show that the chromosome segregation defect is exacerbated by Ter DNA amplification. Indeed, overproducing topo IB or RNase HI (22) to reduce Ter DNA amplification, deleting *tus* (JB260) to avoid it, or displacing the amplified DNA region outside of Ter (JB136) considerably improves the growth and corrects the cellular filamentation defect of *topA topB* null mutants. Thus, we predict that in *topA topB* null cells the incessant replication in the Ter region delays chromosome segregation and growth by overloading the decatenation capacity of topo IV that needs to remove all the interlinks between the daughter chromosomes, up to the last one, before chromosome segregation can occur. To explain the effect of RNase HI or topo IB on Ter DNA amplification, we envision two mechanisms.

The first one is that replication from weak R-loop-dependent origin(s) in the Ter region, triggers DNA amplification that is initiated following the encounter of replication forks with Ter/Tus barriers. The second mechanism is based on two recent observations showing that delays in replication (67) or the occurrence of DnaA-independent replication (51), by increasing the time that replication forks are trapped at Ter/Tus barriers, stimulate DNA amplification. In this context, R-loops may contribute to amplification by acting as replication impediments or as origins for DnaA-independent replication. Whatever the mechanism(s) at play, our data show that R-loops stimulate Ter DNA amplification that exacerbates the chromosome segregation defect of *topA topB* null cells.

### NER-dependent replication impairment in *topA topB* null mutants

Our results have revealed that the requirement for a minimal level of topo IV overproduction for the survival of *topA topB* null mutants is related to NER activity that somehow impairs replication fork progression. Interestingly, R-loops have been shown to be cleaved by the NER system in eukaryotic cells (68). This cleavage happens despite the fact the size of the R-loop is much larger than the typical lesions processed by the NER system. Cleavage of the R-loop by the eukaryotic NER system only occurs in the frame of the transcription-coupled repair pathway (TCR) and this unconventional processing has been shown to generate DNA double-strand breaks (DSBs) (69–75). Although the proteins from the bacterial NER system are different from those of the eukaryotic system, the recognition and processing mechanism(s) of both systems are similar. Interestingly, a link between R-loop formation and the NER-system acting with Mfd (TCR) in *E. coli* has been recently demonstrated and proposed to involve head-on (HO) transcription-replication conflicts (76). In this case, the conflict in UV-irradiated *rnhA rnhB* (RNase HII) null cells involving a transcription complex arrested by a pyrimidine dimer with a replication fork was hypothesized to trigger R-loop formation. R-loops were shown to interfere with chromosome synthesis that was also found to require replication restart for its completion.

It is possible that in *topA topB* null mutants the processing of R-loops by the NER system can also generate DSBs. The repair of these DSBs involves the action of RecA for the formation of a D-loop that is used by the primosome for replication restart (60). This step would be retarded in the strain with a low level of primosome activity as is the case for the *topA topB* null mutant carrying the *dnaT18::aph* mutation. This could certainly affect replication fork progression as observed in strain JB325. Another possibility is that there is a delay in the resolution of recombination intermediates (e.g., Holliday junctions) that accumulate in this situation. The replication in the presence of a high level of unprocessed recombination intermediates would generate a topological stress (38) that could be resolved by topo IV overproduction.

HO conflicts in *E. coli* can be avoided by *rpoB* mutations conferring a stringent-like phenotype to RNAP (61,77,78). That HO conflicts occur in the absence of topo I is supported by our previous finding that an *rpoB* rifampicin-resistant mutation isolated in a *topA* null mutant that confers a stringent-like phenotype to RNAP, improved the growth of this *topA* mutant (79). Moreover, it was also shown that this *topA rpoB* mutant required less RNase HI activity for optimal growth and accumulated less R-loop-dependent hyper-negatively supercoiled DNA. Very recently, it was shown that the well characterized mutation in term of HO conflicts, *rpoB*35* (78), could allow a MG1655 *topA* mutant to survive (80). However, it was obvious that larger colonies of MG1655 *topA rpoB*35*, that likely indicated the accumulation of *parC parE* amplifications in this MG1655 genetic background (see VS111 strain in this work), were generated at a high frequency on LB plates. Moreover, how this *rpoB*35* mutation affected R-loop-dependent hypernegative supercoiling as well as R-loop formation was not tested in this study. Nevertheless, these findings with *rpoB* mutations together with our results showing the presence of a high level of DnaA-independent replication in *topA* mutants including the MG1655 *topA* mutant VS111, strongly suggest the occurrence of HO conflicts in the absence of topo I.

HO conflicts may involve the action of Mfd (81) recruiting the NER system that cleaves the R-loop, leading to the generation of DSBs. Remarkably, Mfd was recently shown to induce the formation of co-transcriptional R-loops by a mechanism involving the topological partitioning of DNA, with one domain being hyper-negatively supercoiled and promoting R-loop formation (82). Type IA topos would relax such transcription-induced negative supercoiling and therefore prevent R-loop formation. Topo IV overproduction by solving the topological stress associated with HO conflicts as shown recently (83) would somehow prevent Mfd-dependent R-loop formation. Since our results of dot-blot experiments failed to demonstrate a significant effect of topo IV overproduction on the accumulation of R-loops, this mechanism of R-loop formation would not be the major one in *topA topB* null cells.

Notwithstanding the mechanism at play, our data support a model in which the action of topo IV or a type IA topo would solve the topological stress preceding or following NER activity. In this context, the *uvrC25* mutation in JB136 and JB336 would allow these strains to survive without topo IV overproduction by inactivating or attenuating the toxic activity of NER on R-loops and/or other structures. This is indeed only survival for JB336 strain carrying the *uvrC* mutation but no *parC parE* amplification as it barely grows. Interestingly, when the NER system is functional and topo IV is not overproduced, our data also reveal a toxic effect of RNase HI overproduction. It is possible that some R-loops, by slowing down replication, prevent DBSs due to replication forks collapse at the site of NER action or, alternatively, that an R-loop is used for replication restart at this site.

### Bacterial Type 1A topos as relaxases and decatenases

Our data strongly suggest that type IA topos act as backups for each other, initially, as relaxases to prevent co-transcriptional R-loop formation that otherwise generates high levels of topological stress, and finally as decatenases to directly deal with this topological stress that also involves the NER system. The important role of type IA topos in decatenation as suggested from the work presented here has been likely underestimated in the previous studies that were performed with *E. coli* cells lacking only topo III (16). In bacteria, such as *Mycobacterium smegmatis*, with only one type IA topo, this enzyme appears to be essential both for relaxation and decatenation (84). Moreover, a major role for a type IA topo in replication termination has been demonstrated in Human mitochondria (85) and in Trypanosoma (86). Evidence for the involvement of a type IA topo in the resolution of HO replication-transcription conflicts and in replication termination has been presented in *S. cerevisiae* (87). We and others have previously proposed that both the process of replication termination where two convergent replication forks meet, and HO conflicts involving replication and transcription complexes, generate similar topological problems (4,88). The high positive supercoiling accumulating in this situation is either relaxed by DNA gyrase or migrates behind the replication forks to generate precatenanes that are removed by the action of topo IV and type IA topos. The same situation may happen when replication forks move toward unresolved recombination intermediates. Thus, our data shows that type IA topos maintain the stability of the genome by preventing, both directly and indirectly, the topological stress mostly originating from R-loops and from the NER system.

## DATA AVAILABILITY

All DNA sequence files used for MFA analysis are available from NCBI (SRA accession number: PRJNA738450). Numerical data for qPCR are provided in Supplementary data.

## SUPPLEMENTARY DATA

Supplementary Data are available at NAR online.

## ACKNOWLEDGMENTS

We thank Michael Wilson (University of Toronto) for the S9.6 antibodies and Fenfei Leng (Florida International University) for *E. coli* RNase HI and strain VS111. We also thank David Kysela and Armelle Le Campion for excellent technical assistance respectively with fluorescence microscopy and flow cytometry, and George Szatmari for English editing. We also acknowledge Gary Leveque from The Canadian Center for Computational Genomics (C3G). CG3 is a Genomics Technology Platform (GTP) supported by the Canadian Government through Genome Canada.

