## supplementary material for "Bacterial type 1A topoisomerases maintain the stability of the genome by preventing and dealing with R-loop-and nucleotide excision repair-dependent topological stress"

**Table S1**. *Escherichia coli* strains and plasmids used.

| Name | Genotype or Relevant Genotype | Reference or Source |
|---|---|---|
| CT77 | RFM443 *ΔtopB::kan* | RFM443 x P1(MD897) |
| CT170 | RFM475 *ΔtopB::kan* | (1) |
| DM800 | *Δ(topA cysB)204 gyrB225 acrA13* | (2) |
| EV1 | VU409 pSK760 | This work |
| EV3 | VU409 pSK762c | This work |
| JB120 | VU410 *ΔyncE::kan* | VU410 x P1(JW1447-1) |
| JB121 | VU410 *ΔyncE* | JB120, *kan* removed by pCP20 |
| JB136 | JB121 *topA20::*Tn*10 IN(1.52-1.84) uvrC25* | JB121 x P1(RFM480) |
| JB137 | VU411 *topA20::*Tn*10* | VU411 x P1(RFM480) |
| JB177 | VU409 *ΔyncE::kan* | VU409 x P1(JW1447-1) |
| JB185 | VU409 *ΔyncE* | JB177, *kan* removed by pCP20 |
| JB187 | VU409 *ΔtusB ::kan* | VU409 x P1(JW1602-3) |
| JB194 | VU409 *ΔtusB* | JB187, *kan* removed by pCP20 |
| JB198 | PS158 pACYC184Δ*tet*5’ | This work |
| JB206 | RFM445 *topA20::*Tn*10* | RFM445 x P1(RFM480) |
| JB208 | JB137 pET11-*parEC* | This work |
| JB217 | VU409 pET11-*parEC* | This work |
| JB219 | JB217 *topA20::*Tn*10* | JB217 x P1(RFM480) |
| JB222 | JB217 *topA20::*Tn*10* | JB217 x P1(RFM480) |
| JB228 | JB198 *ΔtopB::kan* | JB198 x P1(CT77) |
| JB232 | JB206 pET11-*parEC* | This work |
| JB244 | JB206 pET1B | This work |
| JB252 | JB137 pET1B | This work |
| JB256 | VU409 pET1B | This work |
| JB260 | JB194 *topA20::*Tn*10* | JB194 x P1(RFM480) |
| JB265 | JB256 *topA20::*Tn*10* | JB256 x P1(RFM480) |
| JB266 | JB256 *topA20::*Tn*10* | JB256 x P1(RFM480) |
| JB294 | JB256 *topA20::*Tn*10* (selected at 30°C) | JB256 x P1(RFM480) |
| JB303 | VS111 *ΔtopB::kan* | JB116 x P1 (JW1752-1) |
| JB305 | DM800 *ΔtopB::kan* | DM800 x P1(MD897) |
| JB325 | VU441 *gyrA21* | This work |
| JB326 | VU441 *gyrA21 uvrB24* | This work |
| JB335 | JB185 *topA20::*Tn*10* | JB185 x P1(RFM480) |
| JB336 | JB185 *topA20::*Tn*10 uvrC25* | JB185 x P1(RFM480) |
| JB350 | JB303 pSK760 | This work |
| JB352 | JB303 pSK762c | This work |
| JB354 | JB305 pSK760 | This work |
| JB356 | JB305 pSK762c | This work |
| JB393 | JB137 pSK760 | This work |
| JB395 | JB137 pSK762c | This work |

| | | |
|---|---|---|
| JB442 | JB326 pSK760 | This work |
| JB446 | JB325 pSK760 | This work |
| JB450 | JB336 pSK760 | This work |
| JB511 | EV1 *topA20::*Tn*10* | EV1 x P1(RFM480) |
| JB512 | EV3 *topA20::*Tn*10* | EV3 x P1(RFM480) |
| JW1447-1 | Δ*yncE744::kan* | (3) |
| JW1602-3 | Δ*tus758::kan* | (3) |
| JW1752-1 | Δ*topB761::kan* | (3) |
| MD897 | DM4100 Δ*topB::kan* | Lab collection |
| MM84 | RFM443 *rnhA::cam* | (4) |
| PS158 | RFM475 *gyrA*$^{L83}$ *zei-723*::Tn*10* | (5) |
| RFM443 | *Δ(codB-lacI)3 rpsL200 galK2*(Oc) *IN(rrnD-rrnE)1 rph-1* | (6) |
| RFM445 | *Δ(codB-lacI)3 rpsL200 galK2*(Oc) *IN(rrnD-rrnE)1 rph-1 gyrB221* (Cou$^r$) *gyrB203*(Ts) | (6) |
| RFM475 | *Δ(codB-lacI)3 rpsL200 galK2*(Oc) *IN(rrnD-rrnE)1 rph-1 gyrB221* (Cou$^r$) *gyrB203*(Ts) *Δ(topA cysB)204* | (6) |
| RFM480 | *Δ(codB-lacI)3, rpsL200, galK2*(Oc), *IN(rrnD-rrnE)1, rph-1 gyrB221* (Cou$^r$) *gyrB203*(Ts) *topA20*::Tn*10* | (6) |
| VS111 | F- *λ- ilvG- rfb- 50 rph-I ΔtopA* | (7) |
| VU243 | CT170 *ΔrecA306 srlR301*::Tn*10* | (8) |
| VU452 | RFM475 *ΔrecA306 srlR301*::Tn*10* | (8) |
| VU409 | RFM445 *ΔtopB* | (8)[a] |
| VU410 | RFM445 *ΔtopB* | (8)[a] |
| VU411 | RFM445 *ΔtopB* | (8)[a] |
| VU441 | RFM445 *ΔtopB dnaT18::aph topA20*::Tn*10* | |
| pACYC184Δ*tet*5’ | deletion of the 5’ portion of the *tetA* gene that inactivates its translation | (9) |
| pSK760 | *rnhA* gene with its own promoter | (6) |
| pSK762c | like pSK760 but *rnhA* is mutated and inactive | (6)<br>(6) |
| pET11-*parEC* | production of an active ParEC fusion protein | (10) |
| pET1B | production of an active human Topo IB protein | Lab collection |

[a]Three clones of the transduction experiment to introduce a *topB* deletion in RFM445.

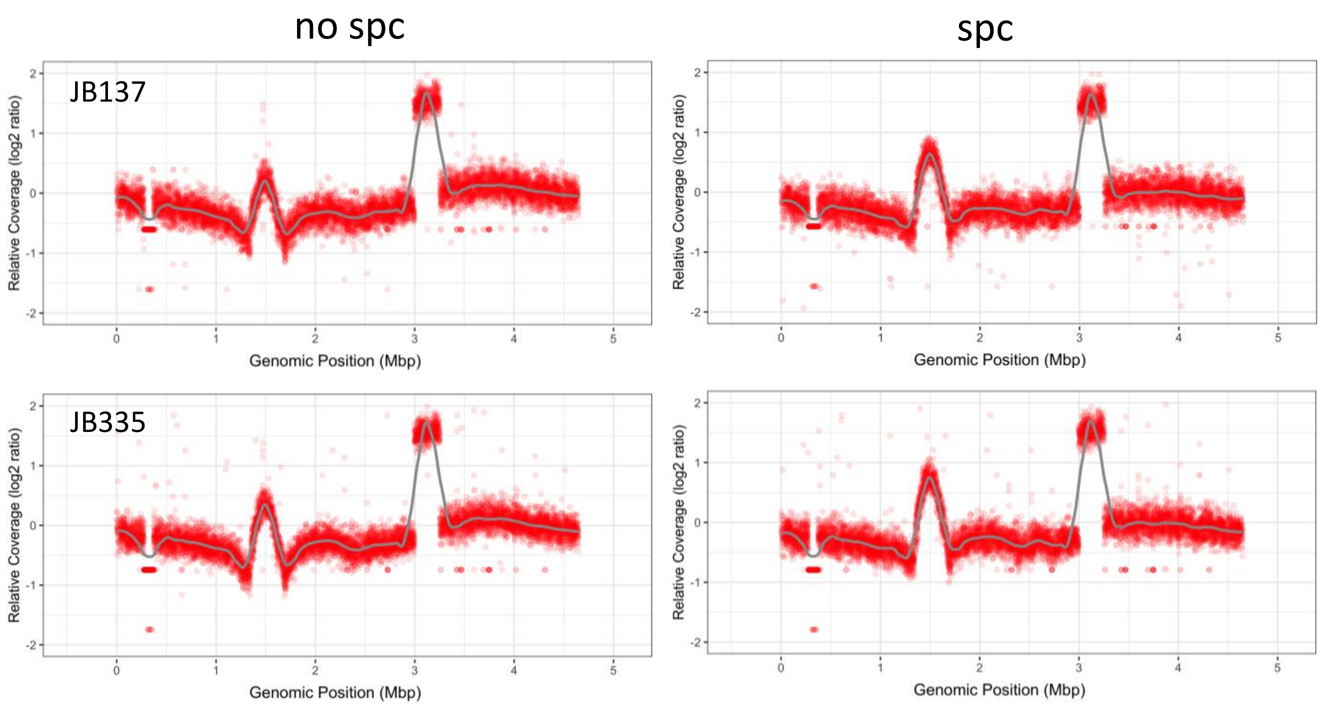


**Figure S1. The *topA topB* null mutants JB137 and JB335 have very similar MFA profiles.** MFA by NGS of genomic DNA extracted from JB137 (*ΔtopB topA20*::Tn*10 gyrB*(Ts)) and JB335 (*ΔtopB topA20*::Tn*10 gyrB*(Ts) *ΔyncE::kan*) cells grown at 30$^{o}$C and treated (spc) or not treated (no spc) with spectinomycin for 2 hours. See legend to Fig. 6 for more details.

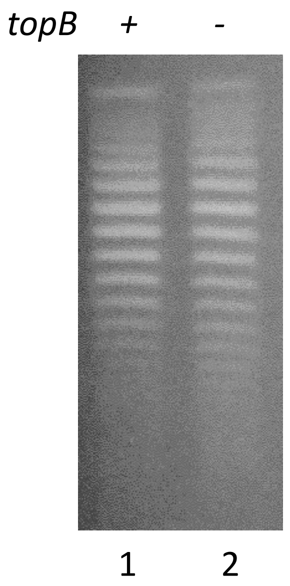


**Figure S2. No effect of a *topB* deletion on the supercoiling level in a *topA*+ *gyrB*(Ts) strain.** One-dimensional chloroquine gel electrophoresis of pACYC184Δ*tet*5' extracted from *gyrB*(Ts) (lane 1) and *gyrB*(Ts) Δ*topB* (lane 2) strains grown at 30°C.

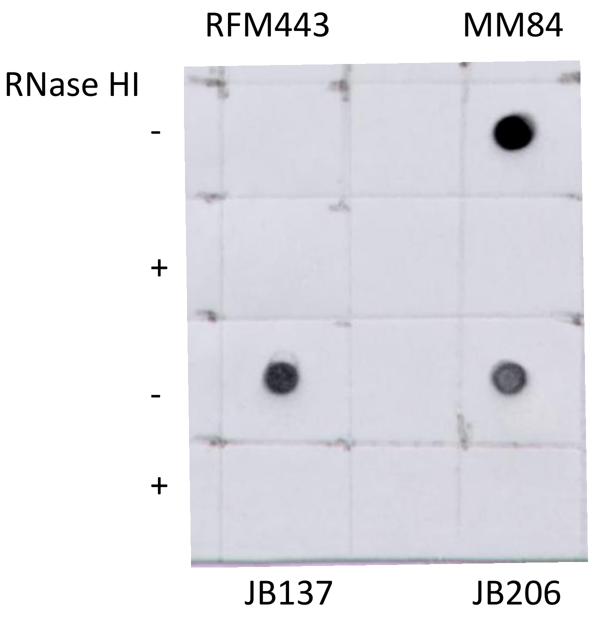


**Figure S3. R-loop formation in type IA topo mutants.** Dot-blot with S9.6 antibodies of genomic DNA from RFM443 (wild-type), MM84 (*rnhA::cam*), JB206 (*topA20*::Tn*10 gyrB*(Ts) and JB137 (*ΔtopB topA20*::Tn*10 gyrB*(Ts)) cells grown at 30$^{o}$C. - and + indicate that DNA was respectively not treated or treated with RNase HI.

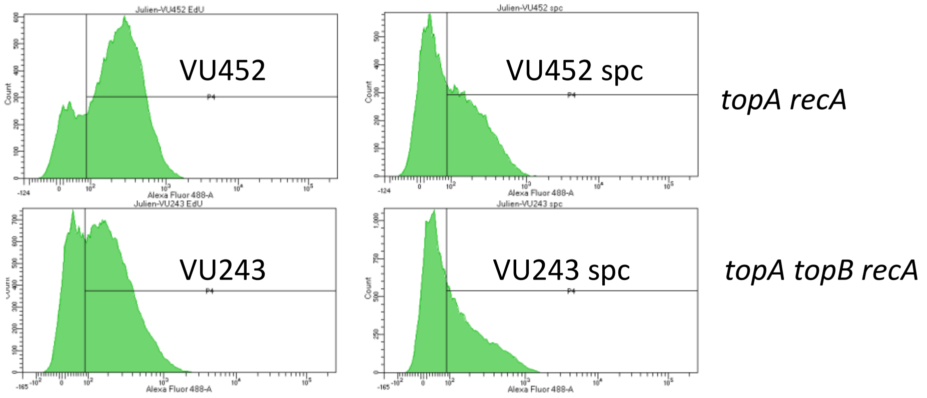


**Figure S4. DnaA-independent replication in *topA* and *topA topB* null mutants is largely *recA*-dependent.** Flow cytometry showing DnaA-independent replication in VU243 (*ΔtopB* Δ*(topA cysB)204 gyrB*(Ts) *ΔrecA306 srlR301*::Tn*10*) and VU452 (Δ*(topA cysB)204 gyrB*(Ts) *ΔrecA306 srlR301*::Tn*10*) cells grown at 30ºC. Spc, means that the cells were treated with spectinomycin for 2 hours before EdU incorporation.

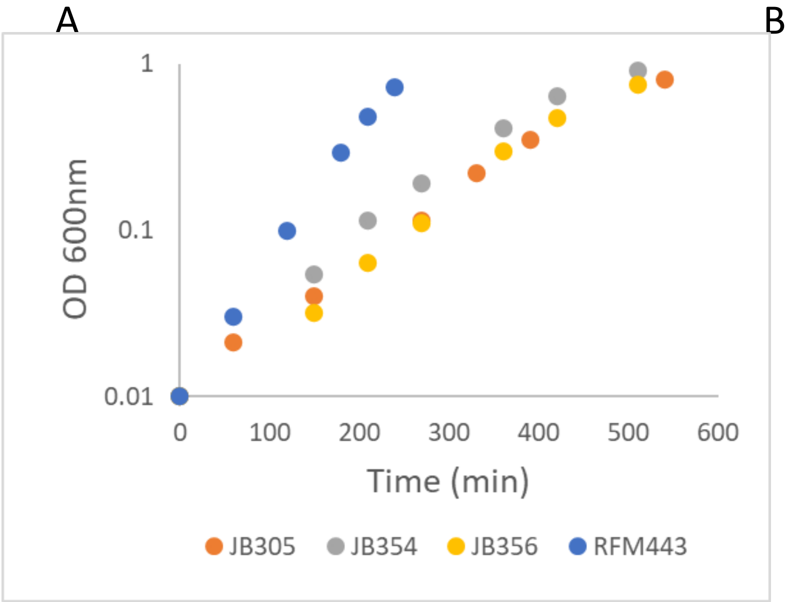


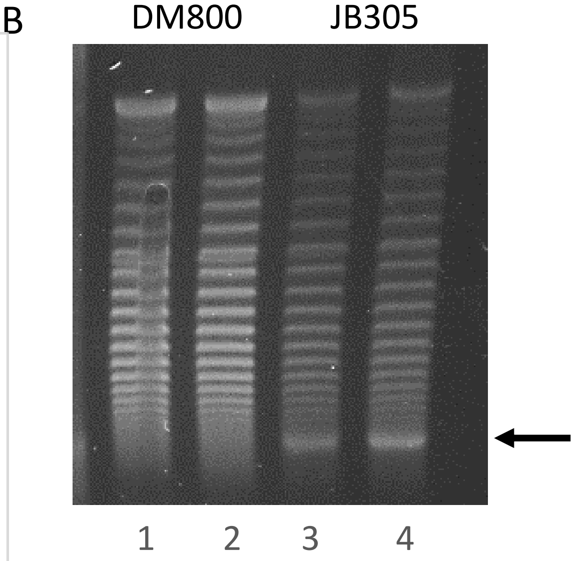


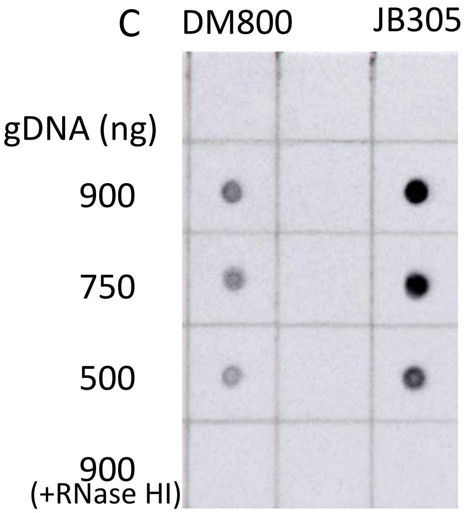


**Figure S5 (continued on next page)**

D

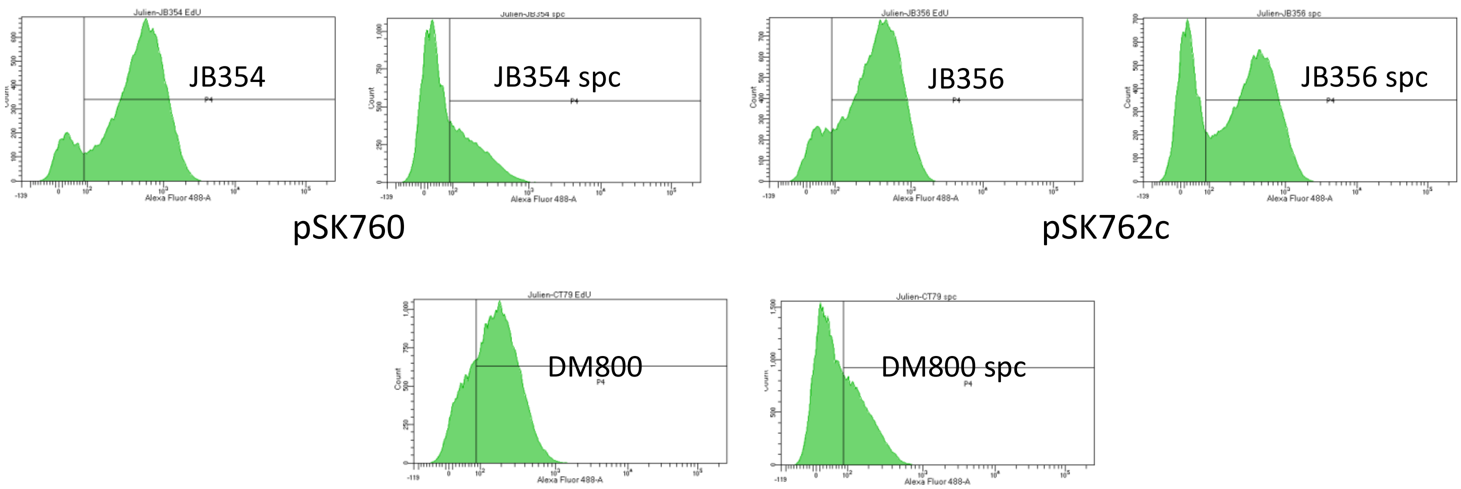


**Figure S5. Phenotypes of a *topB* null derivative of the *topA* null mutant DM800.** (A) Growth curves of RFM443 (wild-type, from Fig. 3A), JB305 (DM800 (*Δ(topA cysB)204*) Δ*topB*), JB354 (JB305/pSK760) and JB356 (JB305/pSK762c) strains at 30$^{o}$C in LB. (B) One-dimensional chloroquine gel electrophoresis of pACYC184Δ*tet5*' extracted from DM800 and JB305 grown at 37$^{o}$C to an OD$_{600}$ of 0.4 (lanes 1 and 3), or 30 min after a transfer from 37 to 30$^{o}$C (lanes 2 and 4). The arrow indicates hyper-negatively supercoiled DNA (C) Dot-blot with S9.6 antibodies of genomic DNA from DM800 and JB305 strains grown at 30$^{o}$C. The amount of genomic DNA spotted on the membrane is indicated. +RNase HI indicates that the genomic DNA was treated with RNase HI. (D) Flow cytometry showing DnaA-independent replication in DM800, JB305, JB354 and JB356 cells grown at 30$^{o}$C. Spc, means that the cells were treated with spectinomycin for 2 hours before EdU incorporation.

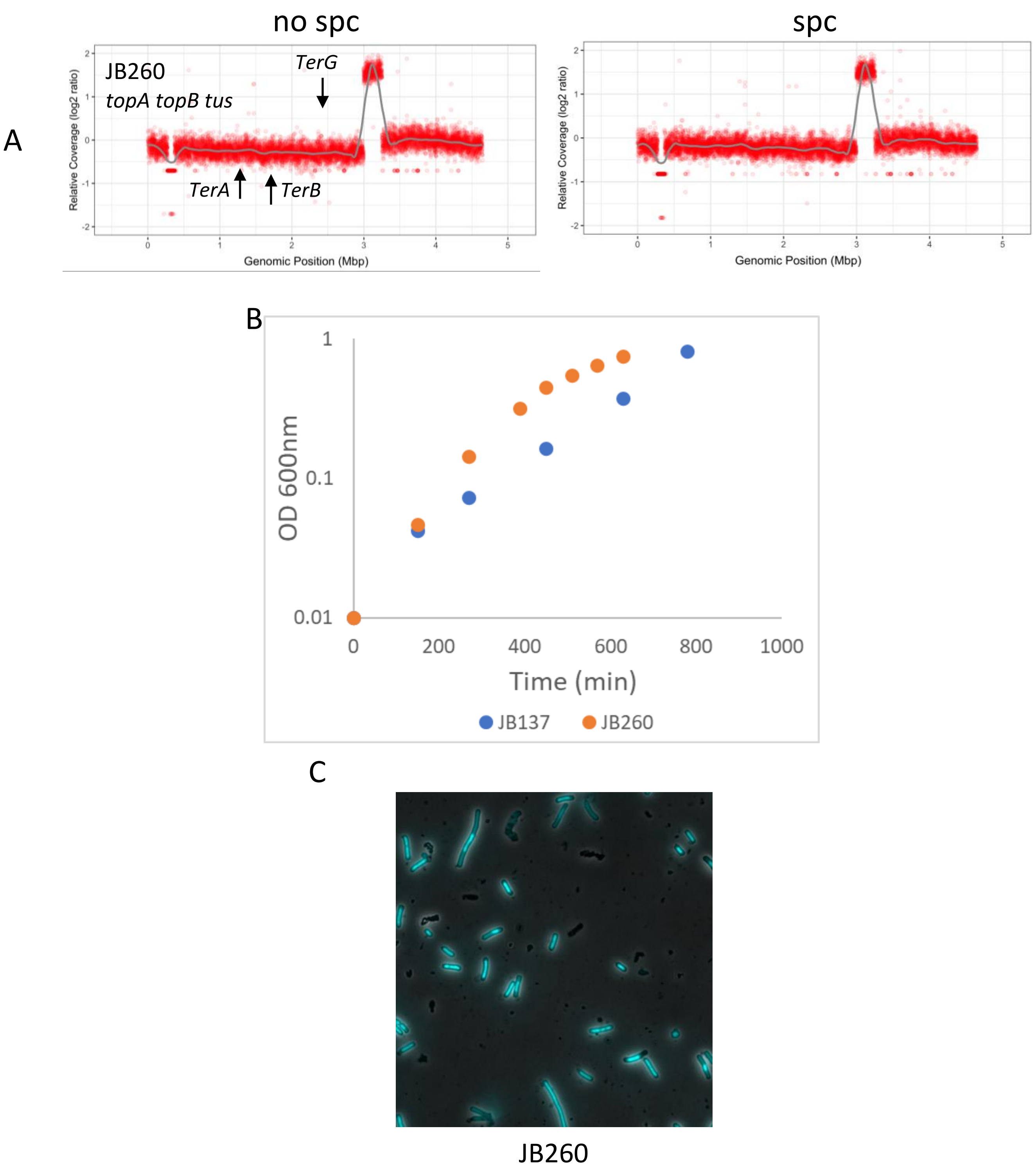


**Figure S6. Effect of a *tus* deletion on phenotypes of a *topA topB* null mutant.** (A) MFA by NGS of genomic DNA extracted from JB260 (Δ*tus ΔtopB topA20*::Tn*10 gyrB*(Ts)) cells grown at 30ºC and treated (spc) or not treated (no spc) with spectinomycin for 2 hours. (B) Growth curves of JB137 (*ΔtopB topA20*::Tn*10 gyrB*(Ts), from Fig. 3A) and JB260 strains at 30ºC in LB. (C) A representative merged image of phase contrast and fluorescence pictures of SYTO-40-stained JB260 cells grown at 30ºC.

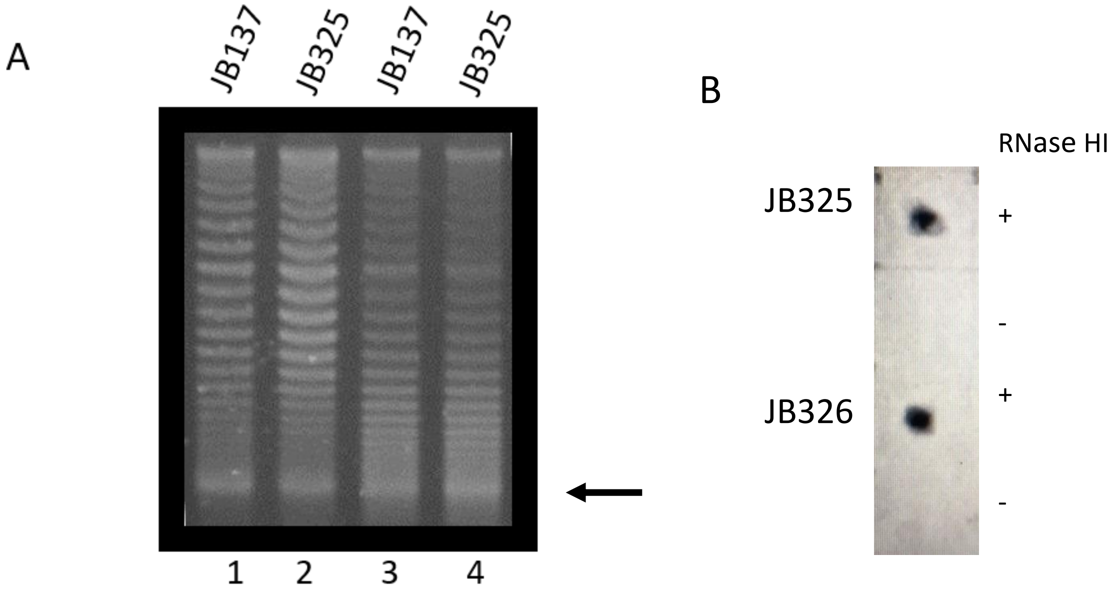


**Figure S7. Hypernegative supercoiling and R-loop formation in *topA topB* null mutants are not affected by *dnaT18::aph, gyrA21* and *uvrB24* mutations.** (A) One-dimensional chloroquine gel electrophoresis of pACYC184Δ*tet5*' extracted from JB137 ((*ΔtopB topA20*::Tn*10 gyrB*(Ts)) and JB3325 (*ΔtopB topA20*::Tn*10 gyrB*(Ts) *gyrA21*) cells grown at 37$^{o}$C to an $OD_{600}$ of 0.4 (lanes 1 and 2), or 30 min after a transfer from 37 to 30$^{o}$C (lanes 3 and 4). (B) Dot-blot with S9.6 antibodies of genomic DNA from JB325 (*ΔtopB topA20*::Tn*10 gyrB*(Ts) *gyrA21*) and JB326 (*ΔtopB topA20*::Tn*10 gyrB*(Ts) *gyrA21*, *uvrB25*) strains grown at 30$^{o}$C. 900 ng of genomic DNA were spotted on the membrane. +RNase HI indicates that the genomic DNA was treated with RNase HI.

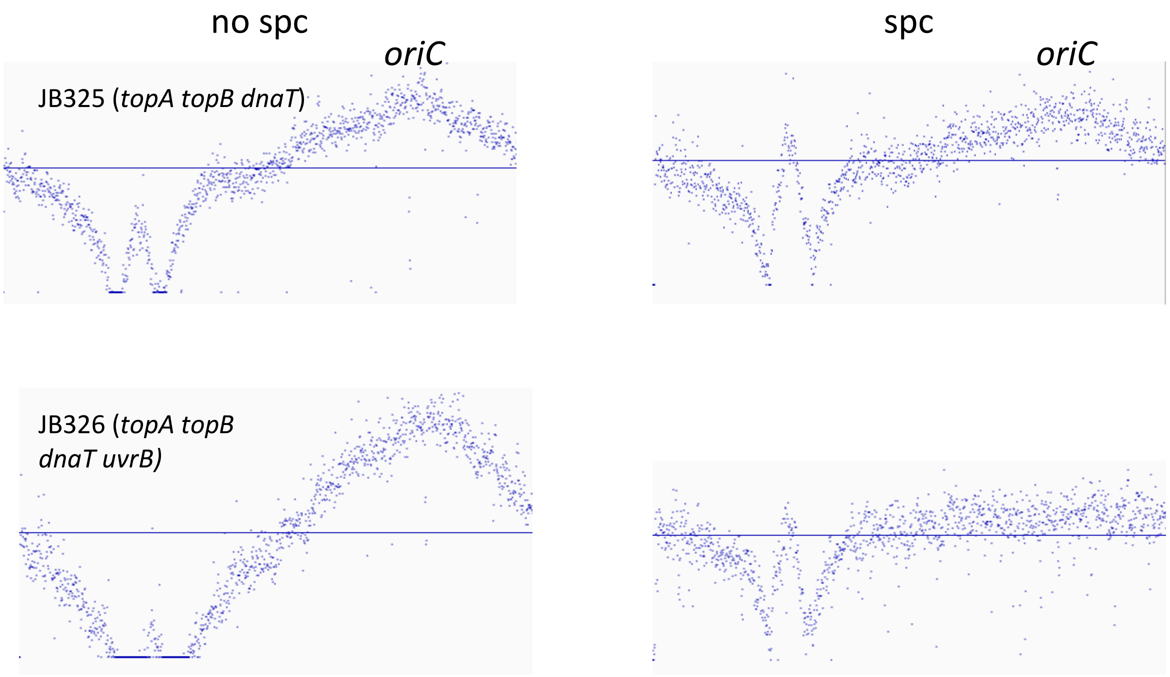


**Figure S8. The effect of the *uvrB24* mutation on the completion of *oriC*-dependent replication in a *topA topB dnaT18::apĥ* mutant not carrying a *parC parE* amplification.** The IGV (Integrative Genomics Viewer) tool was used to generate MFA profiles from the NGS data for strains JB325 (*ΔtopB topA20*::Tn*10 gyrB*(Ts) *gyrA21*) and JB326 (*ΔtopB topA20*::Tn*10 gyrB*(Ts) *gyrA21*, *uvrB25*). It is quite clear here that despite the higher level of replication originating from *oriC* in strain JB326 as compared to strains JB325 (no spc profiles), only in JB326 that replication from *oriC* is fully completed (JB326 *spc*, flat profile).

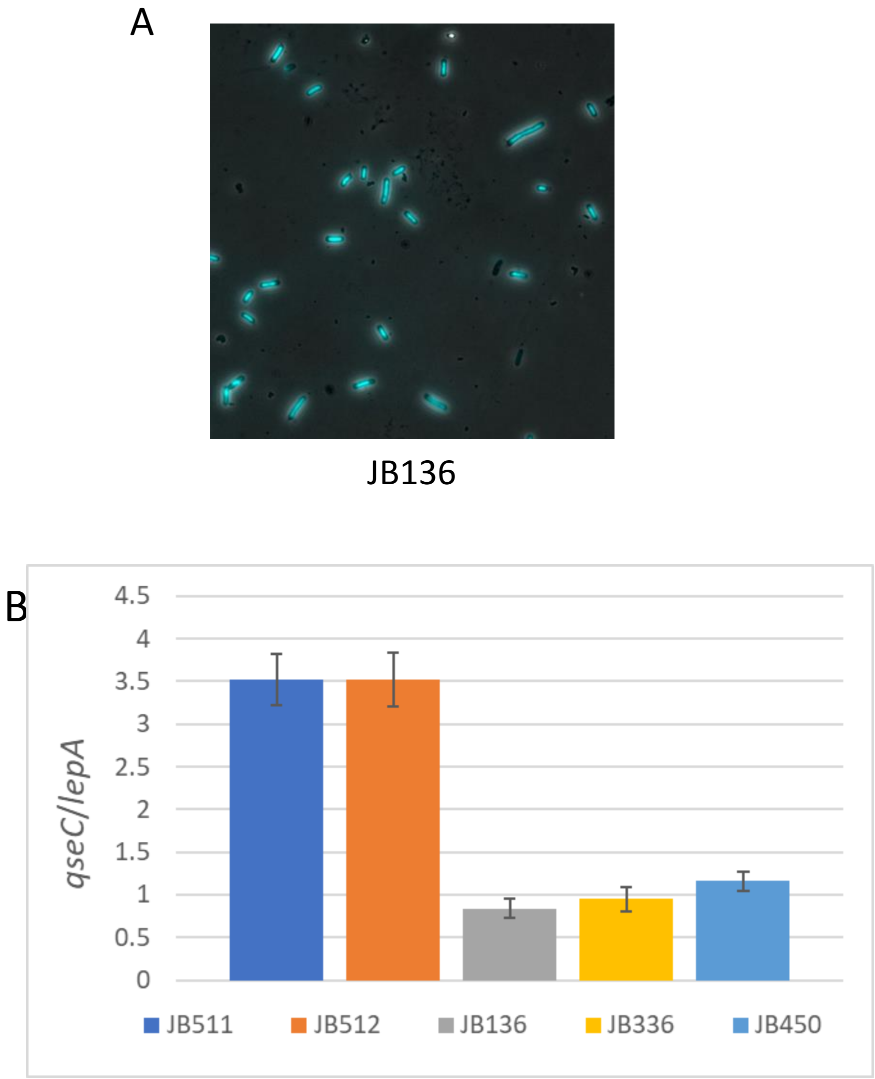


**Figure S9. (A) Cell morphology of a *topA topB* null mutant carrying an inversion involving a portion of the chromosomal Ter region.** A representative merged image of phase contrast and fluorescence pictures of SYTO-40-stained JB136 (*ΔtopB topA20*::Tn*10 gyrB*(Ts), *IN(1.52-1.84)*, *uvrC25*) cells grown at 30ºC. **(B) A *topA* null transductant of *topB* null cells overproducing RNase HI requires *parC parE* amplification for survival unless the *uvrC25* mutation is present.** *qseC/lepA* ratio determined by qPCR of genomic DNA from strains JB136 (*ΔtopB topA20*::Tn*10 gyrB*(Ts), *IN(1.52-1.84)*, *uvrC25*), JB336 (*ΔtopB topA20*::Tn*10 gyrB*(Ts) *uvrC25*), JB450 (JB336/pSK760), JB511 (*ΔtopB topA20*::Tn*10 gyrB*(Ts)/pSK760) and JB512 (*ΔtopB topA20*::Tn*10 gyrB*(Ts)/pSK762c).
